# Tubulin glutamylation is key to axon guidance via selective tuning of microtubule-severing enzymes

**DOI:** 10.1101/2022.01.20.477127

**Authors:** Daniel Ten Martin, Nicolas Jardin, Juliette Vougny, François Giudicelli, Laïla Gasmi, Véronique Henriot, Laura Lebrun, Cécile Haumaître, Matthias Kneussel, Xavier Nicol, Carsten Janke, Maria Magiera, Jamilé Hazan, Coralie Fassier

## Abstract

The microtubule cytoskeleton is a major driving force of neuronal circuit development. Fine-tuned remodelling of this network by selective activation of microtubule-regulating proteins, including microtubule severers, emerged as a central process in neuronal wiring. Tubulin posttranslational modifications control both microtubule properties and the activities of their interacting proteins. However, whether and how tubulin posttranslational modifications may contribute to neuronal connectivity has not yet been addressed. During zebrafish embryogenesis, we show that the microtubule severers p60-katanin and spastin play specific roles in axon guidance and identify a key role for tubulin polyglutamylation in their functional specificity. Furthermore, our work reveals that polyglutamylases with undistinguishable activities *in vitro*, TTLL6 and TTLL11, play exclusive roles in axon navigation by selectively tuning p60-katanin and spastin activities. We confirm the selectivity of TTLL11 towards spastin activation in mammalian cortical neurons and establish its relevance in preventing axonal degeneration triggered by spastin haploinsufficiency. Our work thus provides mechanistic insight on the control of microtubule-driven neuronal development and homeostasis, and opens novel avenues for developing therapeutic strategies in spastin-associated hereditary spastic paraplegia.

## Introduction

Over the past decades, microtubules (MTs) have emerged as key players in nervous system development. In addition to providing mechanical support, MTs form railways for axonal transport and mediate key signalling events that contribute to neuron morphological and behavioural changes during neural circuit wiring (Conde and Caceres, 2009; Hoogenraad and Bradke, 2009; Dent et al., 2011). Numerous studies have established MT remodelling as a key driving force in axon outgrowth and guidance (Sanchez-Huertas and Herrera, 2022; Atkins et al., 2023). Notably, the asymmetric invasion and stabilisation of MTs within filopodia drive directed axon outgrowth (Williamson et al., 1996; Zhou et al., 2002; Buck and Zheng, 2002). Moreover, several guidance cues influence MT dynamics within the growth cone and thereby growth directionality *in vitro*, while a growing number of MT-interacting proteins are required for neuronal circuit wiring *in vivo* (Sanchez-Huertas and Herrera, 2022; Atkins et al., 2023). While these data illustrate MT central role in shaping neuronal connectivity, the intrinsic mechanisms underpinning the fine-tuned remodelling of this network in developing axons are far from being understood.

Importantly, tubulin posttranslational modifications (PTMs) generate sub-populations of MTs that exhibit distinct physical properties and may locally fulfil specific cellular functions (Janke and Magiera 2020; McKenna et al., 2023). Tubulin polyglutamylation, which consists in the addition of glutamate side chains to tubulin C-terminal tails, is an abundant tubulin PTM in neurons (Audebert et al., 1994). This modification is catalyzed by enzymes from the Tubulin Tyrosine Ligase Like family (TTLL; van Dijk et al., 2007). Each TTLL has an enzymatic specificity for either short or long chains of glutamates, and a substrate preference for α- versus β-tubulin (van Dijk et al., 2007). Lately, tubulin polyglutamylation was shown to fine-tune the binding affinity and/or activity of several MT-interacting proteins (Valenstein et al., 2016; Genova et al., 2023) and the dysregulation of its physiological levels impairs neuronal homeostasis in mice and humans (Magiera et al., 2018; Shashi et al., 2018). Yet, the physiological role of this tubulin PTM, as well as the selectivity of TTLL-generated polyglutamylation patterns towards MT-interacting proteins in developing neurons, remains poorly characterised.

Interestingly, TTLL-mediated tubulin polyglutamylation promotes MT severing by spastin and p60-katanin *in vitro* (Lacroix et al., 2010; Valenstein and Roll-Mecak., 2016; Shin et al., 2019; Genova et al., 2023). Notably, these critical regulators of MT mass and organisation (McNally and Roll-Mecak, 2018, Vemu et al; 2018, Kuo et al., 2019) are central for nervous system development and homeostasis (Lynn et al., 2021; Costa and Sousa, 2022) and involved in human cortical malformations for p60-katanin (Eom et al., 2014; Hu et al., 2014; Mishra-Gorur et al., 2014) and hereditary spastic paraplegia for spastin (Hazan et al., 1999). During development, both spastin and katanin play a decisive role in vertebrate axon outgrowth and branching (Ahmad et al., 1999, Karabay et al., 2004; Yu et al., 2008, Butler et al., 2010), while only spastin has been involved in axon guidance to date (Jardin et al., 2018).

We here explore whether TTLL-mediated MT polyglutamylation may constitute an original mechanism controlling neuronal circuit wiring via the selective regulation of severing-enzyme activities. By combining loss-of-function and rescue experiments in zebrafish larvae, we first identify p60-katanin as a novel key player in motor axon navigation and characterise its non-redundant role with spastin in this process. We next provide the first *in vivo* evidence that long chain glutamylases TTLL6 and TTLL11 selectively tune the respective activity of p60-katanin and spastin to control motor axon targeting during development. We further provide a proof of concept that MT polyglutamylation generated by specific TTLL enzymes rescues the axon phenotypes caused by defective spastin gene dosage both in zebrafish larvae and mammalian neurons. Overall, our work pinpoints tubulin polyglutamylation as a central regulator of MT functions in the developing brain as well as an appealing therapeutic target for neurological disorders involving MT-severing enzymes.

## Results

### *p60-Katanin* controls zebrafish spinal motor axon pathfinding and larval locomotion

The established role of MT-severing Spastin in zebrafish motor circuit wiring (Jardin et al., 2018) prompted us to explore its functional similarity/diversity with another MT severer present in developing axons, p60-Katanin (Karabay et al., 2004), via loss-of-function analyses. Using *in toto* immunohistochemistry and *in vivo* live imaging approaches in Tg(*Hb9*:GFP) transgenic larvae expressing GFP in motor neurons, we showed that *p60-katanin* knockdown affected secondary motor axon (sMN) targeting in a dose-dependent manner (Fig. 1A-E). Indeed, while low doses of p60-Katanin morpholino (MO^p60Kat/1.3^ ^pmol^) led to the abnormal split and targeting of the dorsal nerve (empty arrows, Fig 1A-B and Movies EV1-2), its higher doses (MO^p60Kat/3.4^ ^pmol^) significantly impaired dorsal nerve formation compared to MO^Ctl^ and MO^p60Kat/1.3^ ^pmol^ embryos (full arrows, Fig 1A-C). MO^p60Kat/3.4^ ^pmol^ morphants and to a lesser extent MO^p60Kat/1.3^ ^pmol^ larvae, also exhibited misrouted rostral nerves that were aberrantly targeted caudally (empty arrowheads, Fig 1A,D), a phenotype barely observed in control larvae (full arrowheads, Fig 1A,D and Movies EV3-4). Moreover, 57% of MO^p60Kat/3.4^ ^pmol^ larvae showed sMN axons that exited the spinal cord at ectopic sorting points (asterisks, Fig 1A,E), compared to 10% in MO^p60Kat/1.3^ ^pmol^ and 3% in MO^Ctl^ larvae. Notably, the sMN defects of MO^p60Kat/1.3^ ^pmol^ and MO^p60Kat/3.4^ ^pmol^ morphant larvae were also observed following the injection of another p60-Katanin morpholino (MO^katna1aug1^; Butler et al. 2010; Fig EV1) and were rescued by injecting proportional doses of human *KATNA1* mRNA (Fig 1A-E), confirming that they were specifically due to a dose-dependent lack of p60-Katanin.

**Figure 1.**
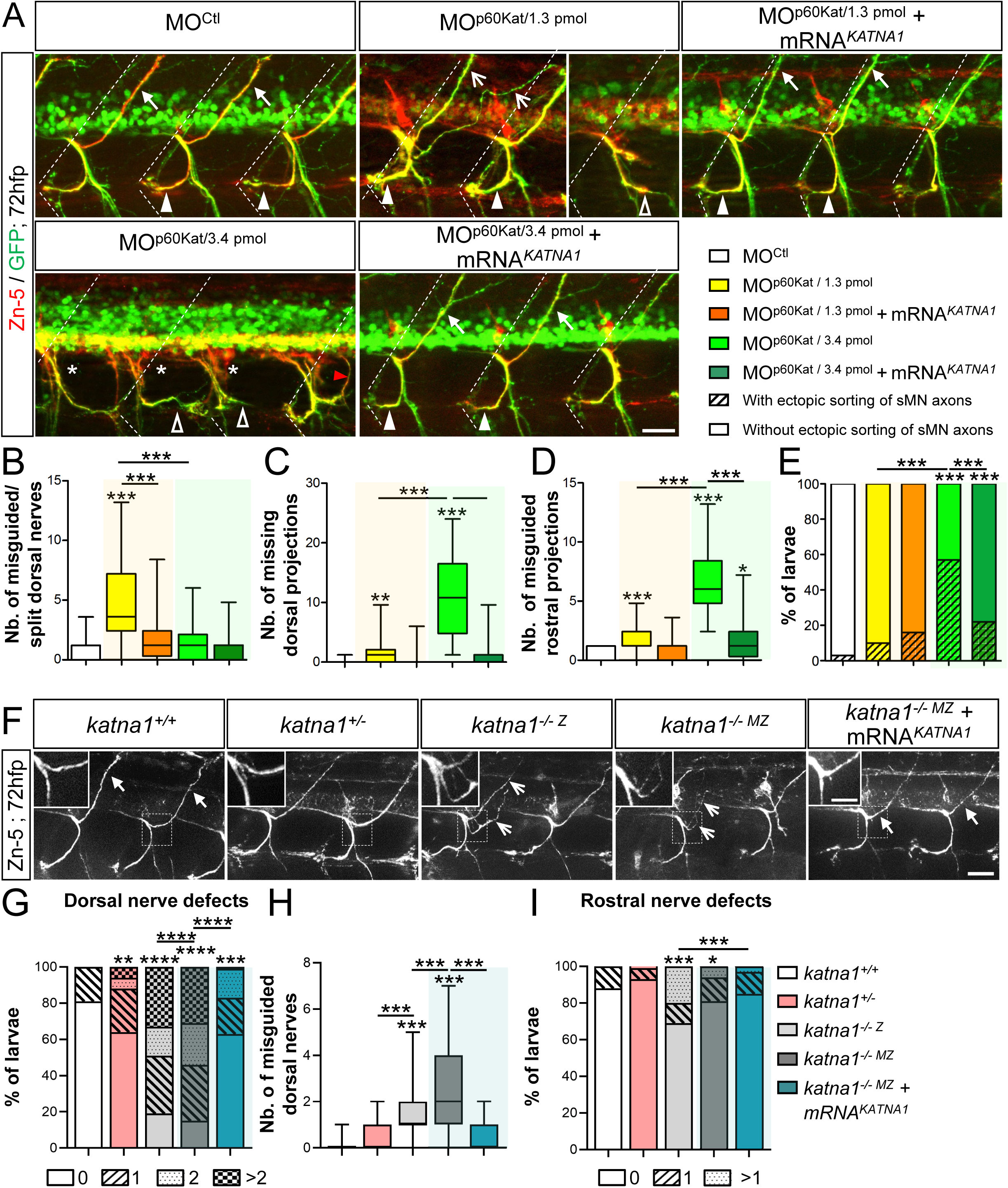
*p60-katanin* impairs the axon pathfinding of secondary motor neurons. (A) Immunolabelling of sMN tracts in 72-hpf Tg(*Hb9*:GFP) larvae injected with control morpholino (MO^Ctl^; n=32), *p60-katanin* morpholino (MO^p60Kat*/*1.3pmol^, n=32 and MO^p60Kat*/*3.4pmol^, n=32) or co-injected with *p60-katanin* morpholino and human *KATNA1* mRNA (MO^p60Kat/1.3pmol^ + mRNA*^KATNA1^*, n=32; MO^p60Kat^*^/^*^3.4pmol^ + mRNA*^KATNA1^*, n=32) using Zn-5 and GFP antibodies. Dotted lines mark lateral myosepta (B-D) Mean number of misguided dorsal projections (B), missing dorsal projections (C) and misguided rostral nerves (D) per larva. (E) Percentage of larvae with ectopic sorting of sMN axons from the spinal cord. (F) Immunolabelling of sMN axons in 72-hpf controls (*katna1^+/+^*, n=34), as well as heterozygous (*katna1^+/−^*, n=53), homozygous zygotic (*katna1^-/-Z^,* n=33) and maternal zygotic (*katna1^-/-MZ^,* n=12) mutant larvae using Zn-5 antibody. Insets are higher magnifications of the dorsal nerve. (G) Percentage of larvae with dorsal nerve defects. (H) Mean number of misguided dorsal projections. (I) Percentage of larvae with dorsal nerve defects. (A, F) Lateral views of the trunk, anterior to the left. Full arrowheads and full arrows show normal rostral and dorsal nerves, respectively. Empty arrowheads and empty arrows indicate misguided rostral and split/misguided dorsal projections, respectively. Asterisks point at aberrant exit points of sMN axons from the spinal cord. Scale bars: 25 µm, inset: 10 µm (F). (B-E, G-I) Quantifications were performed on 24 spinal hemisegments around the yolk tube per larva. (B-D, H) Box and Whisker graphs. *p≤ 0.05; **p≤ 0.01; ***p≤ 0.001; Kruskal–Wallis ANOVA test with Dunn’s post hoc test. Whiskers indicate the Min/Max values. (E, G, I) **p≤0.01; ***p≤0.001; Chi^2^ test.

To further validate the specificity of the p60-Katanin morphant phenotype, we analysed the development of sMN axons in the *katna1^sa18250^* mutant (hereafter called *katna1* mutant; Fig 1F-I), harbouring a point mutation (G>A) in the donor splice site of *katna1* intron 4 (Fig EV2A). This mutation caused aberrant insertions of intron-4 fragments into *katna1* transcripts, which all led to a frameshift and the appearance of a premature STOP codon depleting a large portion of the protein including the ATPase domain (Fig EV2B-D). Notably, both zygotic (*katna1^-/-^*^Z^) and maternal zygotic (*katna1^-/-M^*^Z^) *katna1* mutants displayed significant sMN axon pathfinding errors, including the abnormal split of the dorsal nerves and less frequently, the caudal misrouting of the rostral nerves (Fig 1F-I). These anomalies were similar to those described for *p60-Katanin* morphants and were rescued by the overexpression of human *KATNA1* mRNA (Fig 1F-I), strengthening their phenotypic specificity. Furthermore, the increased frequency of dorsal and rostral nerve defects in *katna1^-/-^*^Z^ compared to *katna^+/−^* larva (Fig 1G-I) confirmed that p60-Katanin gene dosage was critical for motor axon targeting. Altogether, these data demonstrate that p60-Katanin controls vertebrate motor axon navigation, in addition to its well-established role in axon extension.

The axon pathfinding defects of *p60-katanin* morphants and mutants (*katna1^-/-MZ^)* were associated with a curved tail phenotype (Movies EV6,8 for morphants and Fig EV2E-F for mutants) as well as striking locomotor deficits in the touch-evoked escape response test at 72 hours post-fertilisation (hpf). These motility defects were characterized by reduced swimming speed and covered distances compared to control larvae (Fig 2A-F and Movies EV5-9) and were rescued by the injection of adequate doses of human *KATNA1* mRNA (Fig 2A-F and Movies EV7,9). These data were consistent with the rescue of the sMN axon guidance defects (Fig 1) and confirmed that these locomotor phenotypes could be specifically assigned to p60-Katanin deficiency. Remarkably, although less severely affected, *katna1* mutants recapitulated the morphological, behavioural and axon pathfinding defects of *p60-katanin* morphants. These results ruled out the possibility of non-specific effects due to morpholino injection or chemical ENU mutagenesis. RT-qPCR analysis of p60-Katanin-related protein-encoding transcripts (Monroe and Hill., 2016) revealed a significant increase in the expression levels of two MT-severing enzymes, Spastin (*spast*), and Fidgetin (*fign*), as well as a *katna1* paralogue, Katanin-like 2 (*katnal2*) in *katna1^-/-M^*^Z^ embryos compared to controls (Fig EV2G). This analysis suggests that genetic compensation by some *katna1*-related genes may account for the milder phenotype of *katna1* null mutants. Altogether, our data establish a specific role for p60-Katanin in zebrafish motor circuit wiring and larval locomotor behaviour.

**Figure 2.**
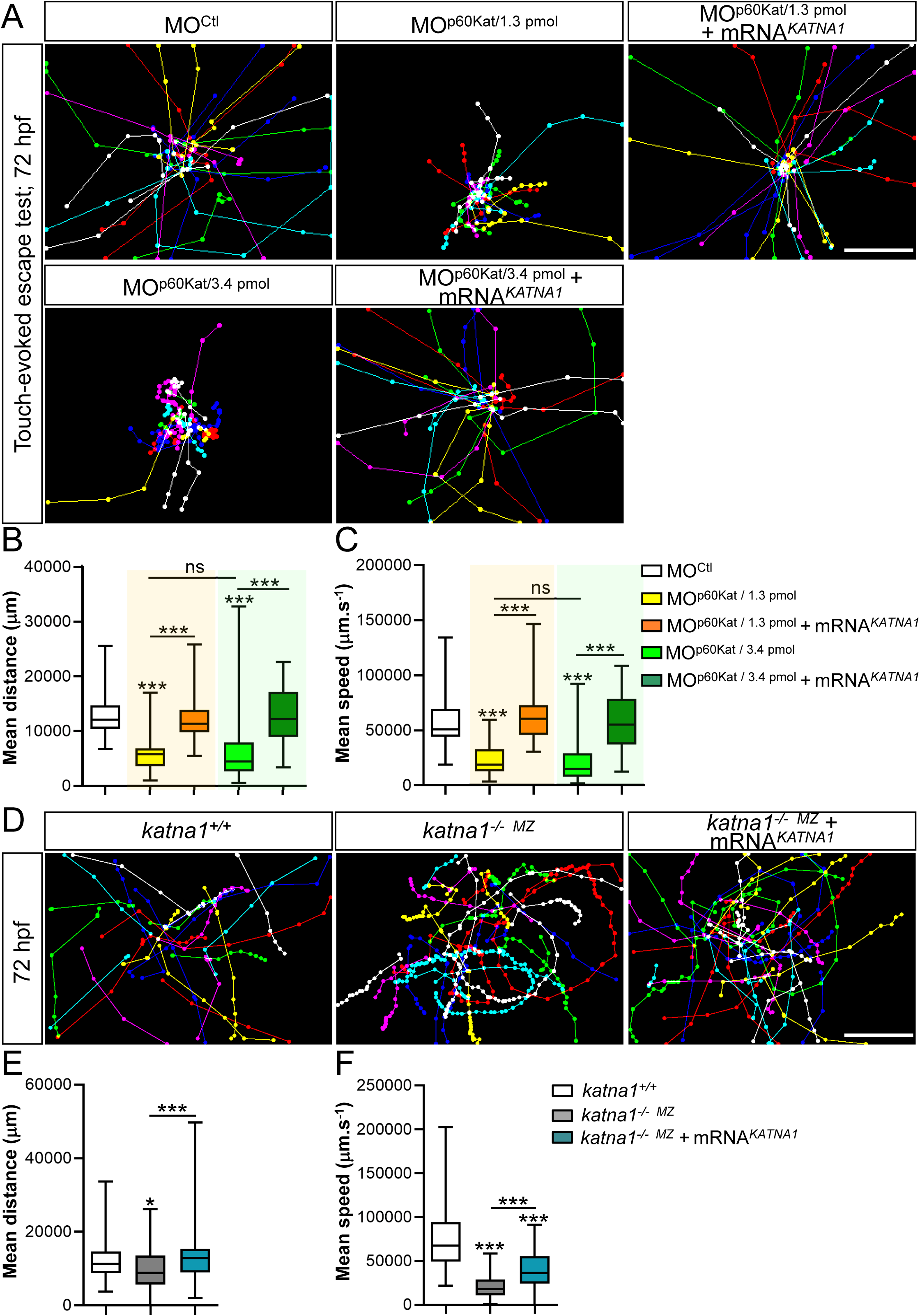
Loss of p60-Katanin causes a dramatic decrease in zebrafish larval mobility. (A) Touch-evoked escape behaviour of 72-hpf larvae injected with a control morpholino (MO^Ctl^; n= 90), increasing doses of *p60-katanin* morpholino (MO^p60Kat/1.3pmol^; n=102 and MO^p60Kat/3.4^ ^pmol^; n=85), or co-injected with MO^p60Kat^ morpholino and human *KATNA1* mRNA (MO^p60Kat/1.3pmol^ + mRNA*^KATNA1^*, n= 91; MO^p60Kat/3.4pmol^ + mRNA*^KATNA1^*, n= 97) (B-C) Mean swimming distance (B) and speed (C) of the larvae tracked in panel A. (D) Touch-evoked escape behaviour of 72-hpf control (*katna1^+/+^*; n=65) and *katna1* mutant larvae injected or not with human *KATNA1* mRNA (*katna1^-/-MZ^;* n=74 and *katna1^-/-MZ^*+ mRNA*^KATNA1^*; n=78). (E-F) Mean swimming distance (E) and speed (F) of the larvae tracked in panel D. (A, D) Each line represents the trajectory of one larva after touch stimulation while the distance between two dots indicates the distance covered by a larva between two consecutive frames. Scale bar: 5mm. (B-C; E-F) Box and Whisker graphs. ***p≤0.001; ns: non-significant (p>0.05); Kruskal–Wallis ANOVA test with Dunn’s post hoc test. Whiskers indicate the Min/Max values.

### P60-Katanin and Spastin play non-overlapping roles in motor circuit wiring

Zebrafish p60-Katanin and Spastin were suggested to have different although related functions in primary motor neuron axon outgrowth (Butler et al., 2010). To support this paradigm, we showed that the depletion of p60-Katanin or Spastin in zebrafish embryos caused eminently different sMN defects, including the abnormal split of the dorsal nerve (p60-Katanin, Fig 1A-B) or the ectopic sorting of motor neuron somata from the spinal cord (Spastin, Jardin et al., 2018). However, both morphants/mutants also exhibited overlapping sMN axon pathfinding defects (e.g., the aberrant caudal targeting of the rostral nerve; Fig 1A,D and Jardin et al., 2018) that could reflect partial functional redundancy. To assess the functional specificity of these closely related MT-severing enzymes in motor neuron development, we conducted cross-rescue analyses. We showed that ubiquitous overexpression of human spastin (i.e., injection of 150 pg/embryo of human spastin-encoding *SPG4* mRNA) failed to rescue the sMN axon pathfinding defects of MO^p60Kat*/*1.3pmol^ morphant (Fig 3A-C) unlike human p60-katanin overexpression (Fig 1), while the same dose of human *SPG4* mRNA efficiently rescued the sMN axon pathfinding errors of zebrafish *spastin* morphants (Jardin et al., 2018). Conversely, ubiquitous overexpression of human p60-katanin (using the same dose of *KATNA1* mRNA as above, see Fig 1) failed to alleviate the axon targeting defects of Spastin-depleted sMN axons (Fig. 3D-F). These data demonstrate that the microtubule-severing enzymes p60-Katanin and Spastin play non-overlapping key roles in vertebrate motor axon navigation.

**Figure 3.**
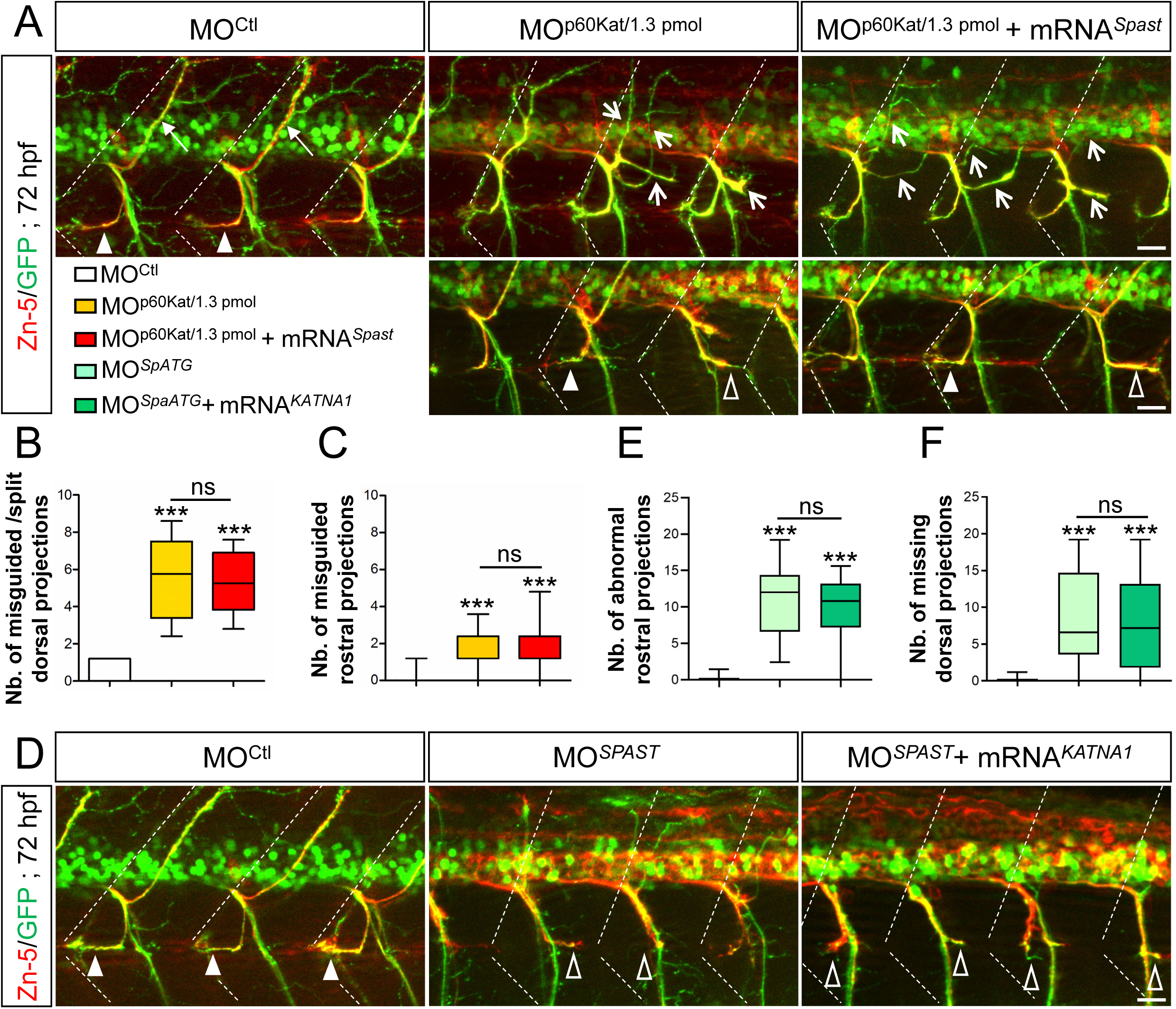
Spastin and p60-katanin have non-redundant roles in motor axon guidance. (A) Immunolabelling of sMN axons in 72-hpf Tg(*Hb9*:GFP) larvae injected with MO^Ctl^ (n= 10), 1.3 pmol of MO^p60Kat^ (n= 10) or co-injected with MO^p60Kat^ and human *SPG4* mRNA (MO^p60Kat/1.3^ ^pmol^ + mRNA*^Spast^*, n= 10) using Zn-5 and GFP antibodies (B-C) Mean number of misguided/split dorsal nerves (B) and misguided rostral nerves (C) per larva. (D) Immunolabelling of sMN axons in 72-hpf Tg(*Hb9*:GFP) larvae injected with MO^Ctl^ (n=29), MO*^spATG^* (n=30) or co-injected with MO*^spATG^* and human *KATNA1* mRNA (MO*^spATG^* + mRNA*^KATNA1^*, n= 29) using Zn-5 and GFP antibodies. (E**-**F) Mean number of abnormal rostral nerves (i.e., caudally targeted or missing; E) and missing dorsal nerves (F) per larva. (A, D) Dotted lines indicate lateral myosepta. Full arrowheads and full arrows show normal rostral and dorsal nerves, respectively. Empty arrowheads and empty arrows indicate misguided rostral and dorsal projections, respectively. All images are lateral views of the trunk, anterior to the left. Scale bars: 25 µm. (B-F) Quantifications were performed on 24 spinal hemisegments around the yolk tube per larva. Box and Whisker graphs. ***p≤ 0.001; ns: non-significant (p>0.05); One-way ANOVA with Bonferroni’s post hoc test (B) and Kruskal–Wallis ANOVA test with Dunn’s post hoc test (C, E-F). Whiskers indicate the Min/Max values.

### The knockdown of tubulin glutamylases TTLL6 and TTLL11 affects motor axon targeting in a similar way to p60-Katanin and Spastin loss of function

Despite their similar expression pattern in the zebrafish developing spinal cord (Fig EV3 for *p60-katanin* and Arribat et al., 2020 for *spastin*), the MT severers p60-Katanin and Spastin have non-redundant functions in motor circuit wiring (Fig 3). This finding prompted us to assess whether the functional specificity of these two enzymes could be underlain by their selective preference for distinct populations of MTs. We focused our analysis on tubulin polyglutamylation, since this posttranslational modification is highly abundant in neurons (Janke and Kneussel, 2010) and was shown to promote spastin and katanin severing activity *in vitro* (Sharma et al., 2007; Lacroix et al., 2010; Valenstein and Roll-Mecak, 2016, Shin et al., 2019, Genova M et al., 2023). Tubulin polyglutamylation is catalysed by Tubulin Tyrosine Ligase-Like (TTLL) enzymes, whose role in neuronal circuit wiring has never been addressed to date. The enrichment of *ttll6* and *ttll11* transcripts in the zebrafish developing spinal cord at stages of motor axon outgrowth (Pathak *et al*., 2011) incited us to examine whether these two long-chain-generating glutamylases could be involved in this developmental process. We first verified the presence of polyglutamylated MTs in zebrafish spinal motor axons using two antibodies recognizing either both long and short chains of glutamates (GT335) or long chains only (polyE). Glutamylation was detected in both pMN axons and sMN nerve tracts at 26 and 72 hpf (Fig 4A-B). We next used a morpholino-based knockdown approach to assess whether TTLL6 and TTLL11 loss-of-function (MO^TTLL6^ or MO^TTLL11^) affected sMN axon pathfinding. While MO^TTLL6^- or MO^TTLL11^-injected 72-hpf larvae were indistinguishable in terms of gross morphology with a ventrally curved body axis characteristic of ciliary mutants (Fig 4C, upper panels), both morphants showed substantially different sMN axon guidance phenotypes (Fig 4C middle and bottom panels). TTLL6-depleted larvae exhibited a significant number of split and misguided dorsal nerves (empty arrows, Fig 4C-D) compared to control larvae (full arrows, Fig 4C-D), as described for MO^p60Kat*/*1.3pmol^ larvae (Fig 1A-B). In contrast, numerous dorsal tracts were missing in *TTLL11* morphants compared to *TTLL6* morphants and controls (Fig 4C,E). Moreover, while 25% and 19% of sMN rostral projections were aberrantly targeted caudally in *TTLL6* and *TTLL11* morphants (empty arrowheads, Fig. 4C,F), the rostral nerves of TTLL11-depleted larvae were mostly defasciculated or missing (bracket, Fig 4C), a phenotype reported for *spastin* mutants (Jardin et al., 2018) and barely observed in *TTLL6* morphant and control larvae (Fig 4C,G). Furthermore, *TTLL11* and, to a lesser extent, *TTLL6* morphants showed sMN axons that exited the spinal cord at ectopic sorting points (asterisks, Fig 4C,H), and ectopic sMN somata outside of the spinal cord (yellow arrows; Fig 4C,I). These two phenotypes were never observed in control larvae (Fig 4C,H-I). Notably, a similar migratory defect of sMN somata misrouted outside of the spinal cord was a characteristic feature of *spastin* morphants and mutants (Jardin et al., 2018).

**Figure 4.**
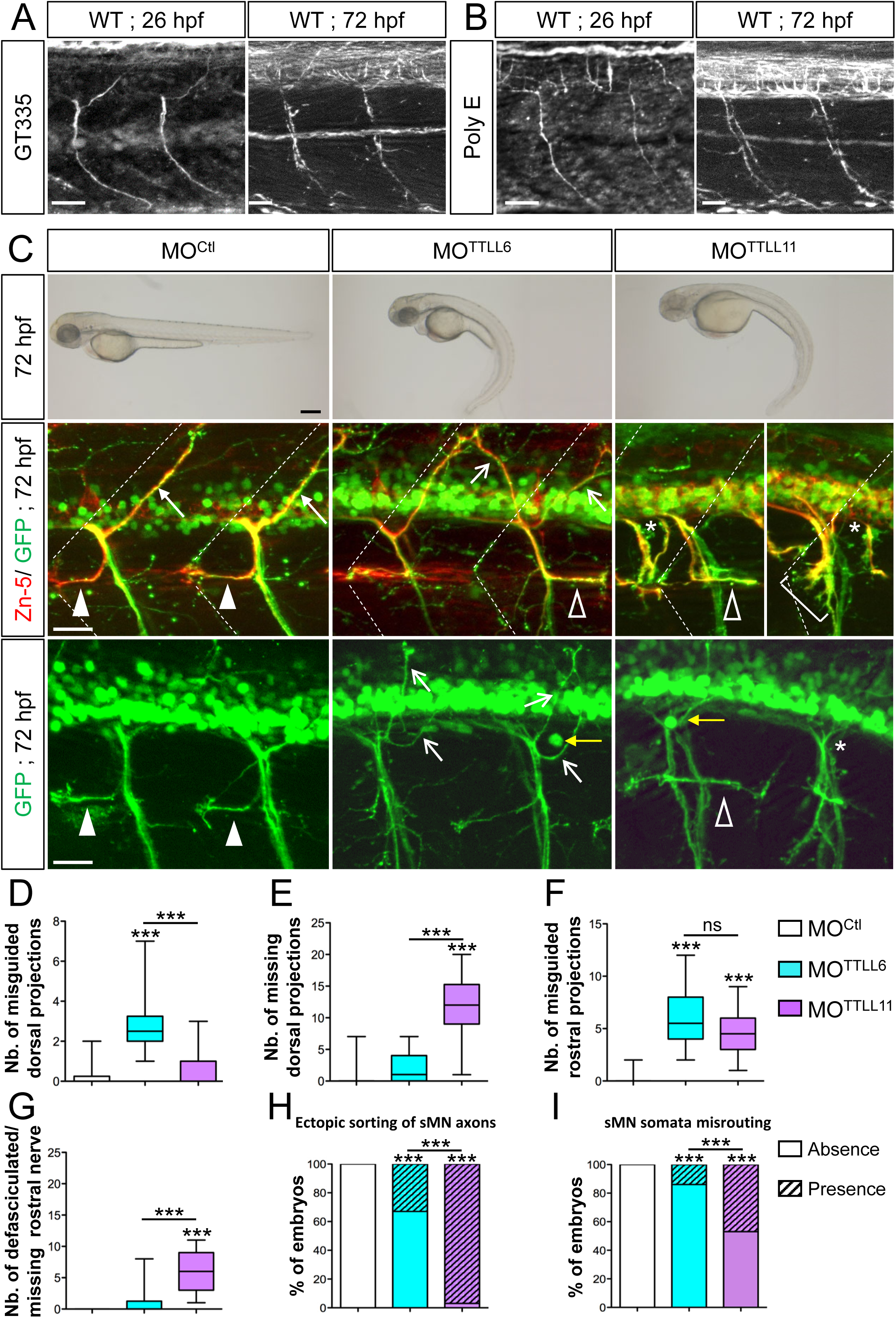
*TTLL6* and *TTLL11* knockdown leads to different motor axon pathfinding defects mimicking the respective phenotypes of p60-Katanin- and Spastin-depleted larvae. (A-B) Immunolabelling of polyglutamylated microtubules (MTs) in 26- and 72-hpf wild-type embryos using GT335 (A) and poly E (B) antibodies. Polyglutamylated MTs are observed in pMN (26 hpf) and sMN (72 hpf) axons. Scale bars: 25 µm. (C) Upper panels: Overall morphology of 72-hpf control (MO^Ctl^), *TTLL6* (MO^TTLL6^) and *TTLL1* (MO^TTLL11^) morphant larvae. Both *TTLL6* and *TTLL11* morphants exhibit a severe ventrally curved body-axis phenotype compared to MO^Ctl^-injected larvae. Scale bars: 250 μm. Middle and bottom panels: Immunolabelling of sMN axons in 72-hpf Tg(*Hb9*:GFP) larvae injected with MO^Ctl^ (n= 30), MO^TTLL6^ (n= 30) or MO^TTLL11^ (n= 30) larvae using Zn-5 and/or GFP antibodies. Dotted lines delineate lateral myosepta. Full arrowheads and full arrows respectively point at normal rostral and dorsal nerves. Empty arrowheads and empty arrows respectively indicate misguided rostral and dorsal projections. Brackets show defasciculated rostral nerves. Asterisks and yellow arrows respectively indicate the ectopic sorting of spinal motor neuron axons and somata from the spinal cord. Scale bars: 25 µm. (D-I) Quantifications of sMN defects in larvae analysed in panel C. Mean number of split/misguided dorsal nerves (D), missing dorsal nerves (E), misrouted rostral nerves (F) and defasciculated/missing rostral nerves (G) per larva. (H-I) Percentage of larvae with ectopic sorting of sMN axons (H) or somata (I) from the spinal cord. (D-I) Quantifications were performed on 24 spinal hemisegments located around the yolk tube per larva. (D-G) Box and Whiskers graphs. ***p≤0.001; ns: non-significant; Kruskal–Wallis ANOVA test with Dunn’s post hoc test. Whiskers indicate the Min/Max values. (H-I) ***p≤0.001. Chi^2^ test.

Although partially overlapping, the motor neuron phenotypes of *TTLL6* and *TTLL11* morphants suggested that these two as yet indistinguishable polyglutamylases could have different functions in motor circuit wiring. Furthermore, the significant phenotypic similarities caused by *TTLL6* or *TTLL11* knockdown and p60-Katanin or Spastin depletion hinted at functional relationship between tubulin glutamylation and MT severing in vertebrate motor axon targeting.

### TTLL6 and TTLL11 have non-redundant functions in motor axon navigation

To assess the specificity of *TTLL6* and *TTLL11* morphant phenotypes and clarify the functional redundancy or specificity of these related enzymes, we carried out rescue and cross-rescue experiments. Notably, overexpression of mouse TTLL6 but not TTLL11 significantly alleviated the morphological and sMN axon guidance errors of *TTLL6* morphants (Fig 5A, C-H). Reciprocally, overexpression of TTLL11 substantially rescued the morphological and sMN axon pathfinding defects of *TTLL11* morphants unlike TTLL6 overexpression (Fig 5B, C-H).

**Figure 5.**
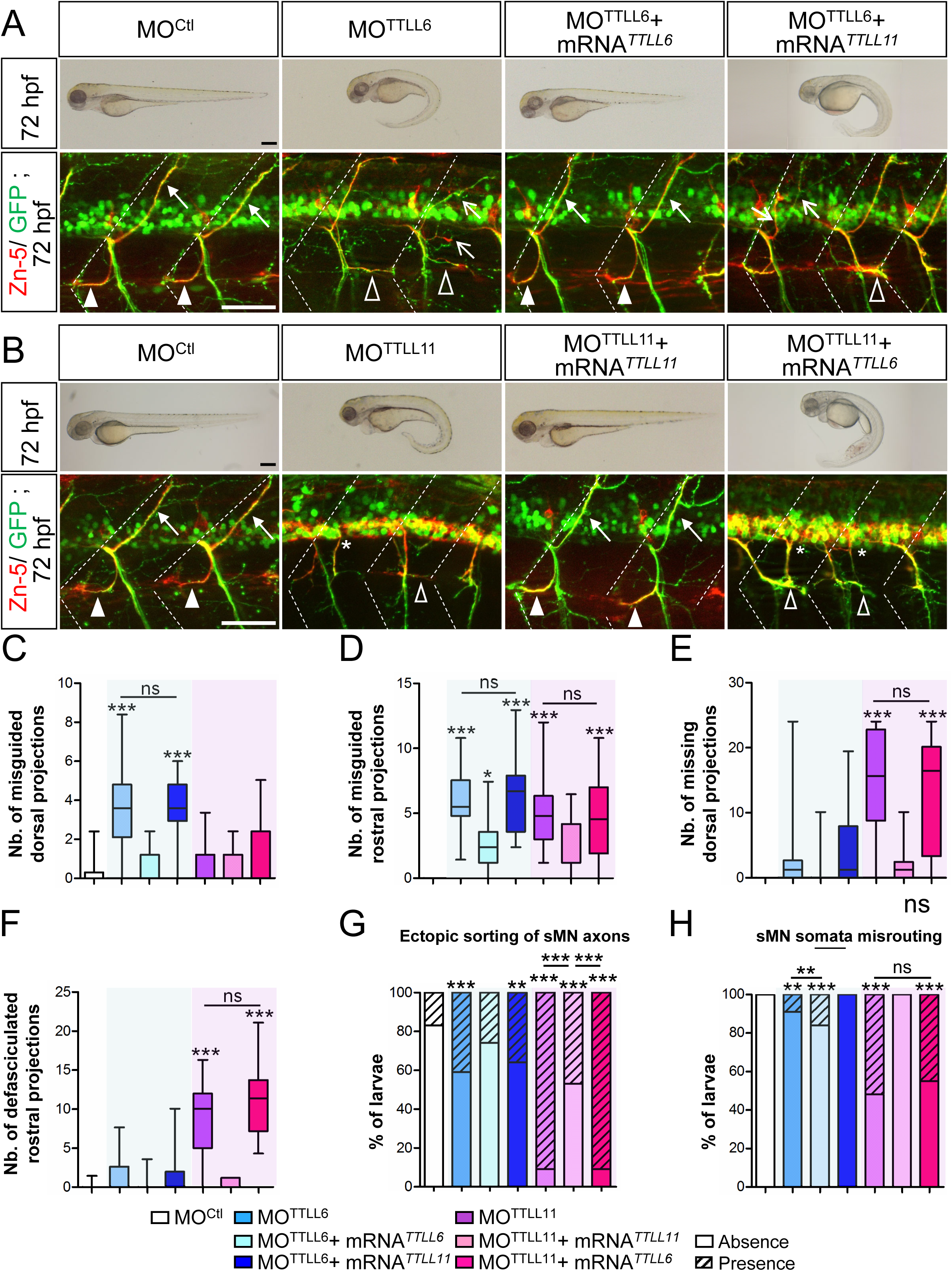
TTLL6 and TTLL11 play non-overlapping roles in spinal motor axon navigation. (A) Rescue and cross-rescue experiments of *TTLL6* morphant phenotypes by co-injection of mouse *TTLL6* or *TTLL11* transcripts. (B) Reciprocal rescue and cross-rescue experiments of *TTLL11* morphant phenotypes by co-injection of mouse *TTLL11* or *TTLL6* transcripts (A-B) Upper panels: Overall morphology of 72-hpf control, morphant and rescued larvae. Scale bars: 250 μm. bottom panels: Immunolabelling of sMN axons with Zn-5 and GFP antibodies in 72-hpf Tg(*Hb9*:GFP) larvae injected with control (MO^Ctl^, n=22), *TTLL6* (MO^TTLL6^, n=22), *TTLL11* (MO^TTLL11^, n=21) morpholinos or co-injected with MO^TTLL6^ or MO^TTLL11^ and mouse *TTLL6* or *TTLL11* mRNA (MO^TTLL6^ *+* mRNA*^TTLL6^*, n=19; MO^TTLL6^ *+* mRNA*^TTLL11^*, n=22; MO^TTLL11^ *+* mRNA*^TTLL11^*, n=17; MO^TTLL11^ *+* mRNA*^TTLL6^*, n=22). Dotted lines delineate lateral myosepta. Full arrowheads and full arrows respectively indicate normal rostral and dorsal nerves. Empty arrowheads and empty arrows respectively point at misguided rostral and dorsal projections. Scale bars: 50 μm. (C-H) Quantifications of sMN defects in larvae analysed in panels A and B. Mean number of split/misguided dorsal nerves (C), misrouted rostral nerves (D), missing dorsal nerves (E) and defasciculated rostral nerves (F) per larva. (G-H) Percentage of larvae with ectopic sorting of sMN axons (G) and somata (H) from the spinal cord. (C-H) Quantifications were performed on 24 spinal hemisegments located around the yolk tube per embryo or larva. (C-F) Box and Whisker graphs. *p≤0.05; ***p≤0.001; ns: non-significant; One-Way ANOVA test with Bonferroni’s post hoc test (D) or Kruskal–Wallis ANOVA test with Dunn’s post hoc test (C, E-F). Whiskers indicate the Min/Max values. (G-H) **p≤0.01; ***p≤0.001; ns: non-significant; Chi^2^ test.

Overall, these analyses reveal that these two glutamylases displaying biochemically similar activities *in vitro* play non-redundant key roles during zebrafish development *in vivo*, and particularly in spinal motor axon navigation.

### Only TTLL6-mediated microtubule polyglutamylation rescues the axon pathfinding and larval locomotor deficit of *p60-Katanin* morphants

The overlapping axon pathfinding phenotypes between *TTLL* morphants and p60-katanin or spastin-depleted larvae led us to hypothesize that these glutamylases could selectively regulate the activity of the two microtubule-severing enzymes during zebrafish motor circuit wiring. To test this hypothesis, we assessed whether promoting TTLL6 or TTLL11-mediated tubulin polyglutamylation could attenuate or rescue the motor axon pathfinding and locomotor deficit caused by p60-Katanin partial depletion by boosting its residual activity. Given that the penetrance of heterozygous *katna1*^+/−^ axon guidance defects is too low to conduct reliable rescue experiments (Fig 1G-H), we addressed the potential rescue effect of TTLL overexpression in *p60-katanin* morphant larvae injected with the lower dose of morpholino, which resulted in robust axon pathfinding defects of the dorsal nerves (i.e., MO^p60Kat/1.3^ ^pmol^; Fig 1A-B). We showed that *TTLL6* but not *TTLL11* overexpression significantly rescued the sMN axon guidance defects (Fig 6A-C) and locomotor deficit (Fig 6A,D-E) of 72-hpf MO^p60Kat*/*1.3pmol^ morphant larvae compared to controls. These results thus suggest that TTLL6 selectively regulates p60-Katanin activity to control motor axon pathfinding and larval locomotion.

**Figure 6.**
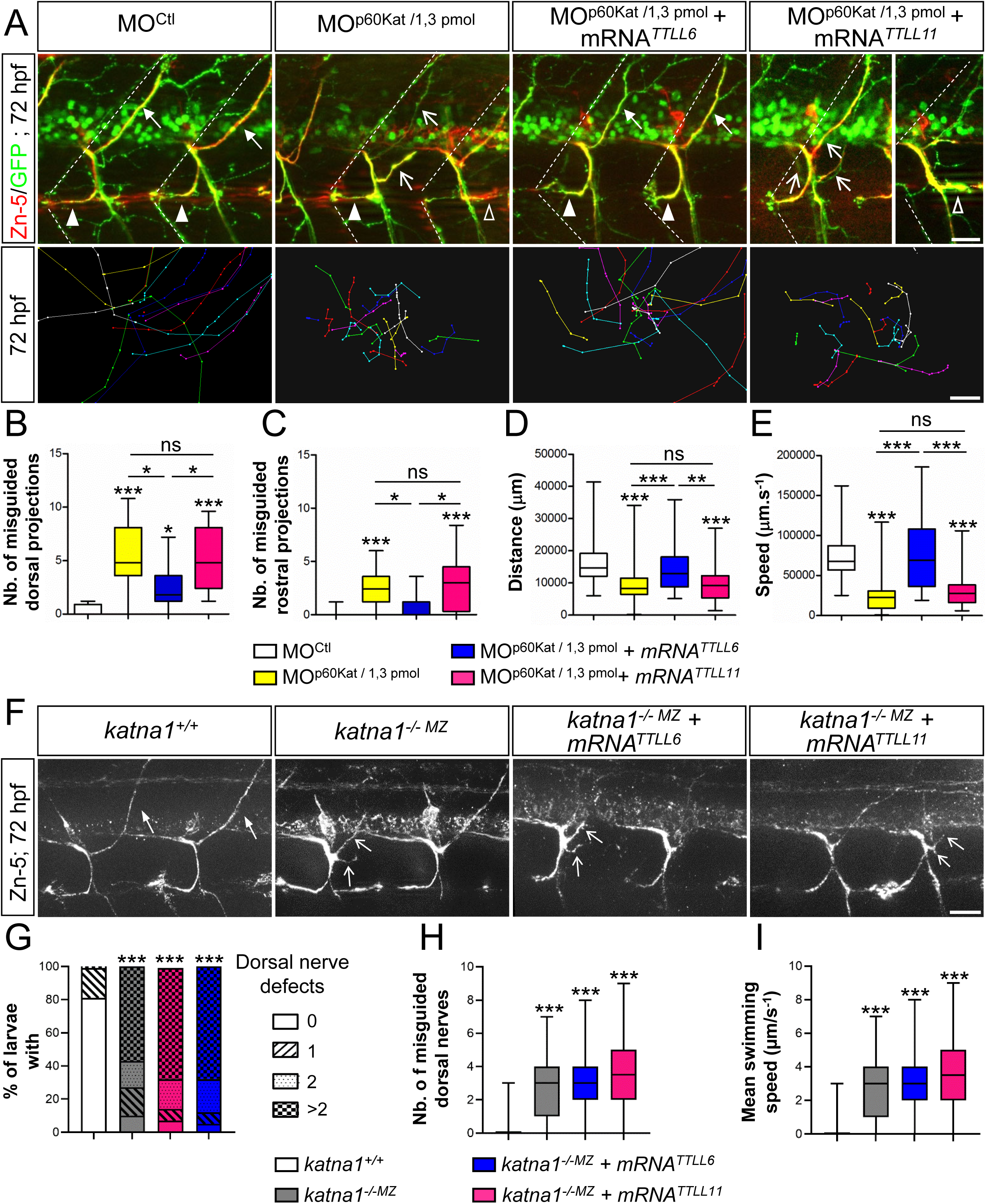
Selective regulation of p60-katanin activity by TTLL6 is required for zebrafish motor axon targeting and larval locomotion. (A) Upper panels: Immunolabelling of sMN axon tracts using Zn-5 and GFP antibodies in 72-hpf Tg(*Hb9*:GFP) larvae injected with MO^Ctl^ (n=20), MO^p60Kat*/*1.3pmol^ (n=20) or co-injected with MO^p60Kat/1.3pmol^ and mouse *TTLL6* (n=19) or *TTLL11* (n=20) mRNAs. Dotted lines delineate lateral myosepta. Lower panels: Tracking analysis of 72-hpf larvae injected with MO^Ctl^ (n=30), MO^p60Kat/1.3pmol^ (n=30) or co-injected with MO^p60Kat/1.3pmol^ and mouse *TTLL6* (n=30) or *TTLL11* (n=30) mRNAs in a touch-evoked escape response test. Each line represents the trajectory of one larva after touch stimulation while the distance between two dots indicates the distance covered by a larva between two consecutive frames. Scale bar: 5 mm (B-E) Quantifications of the sMN and locomotor defects of larvae analysed in panel A. Mean number of split/misguided dorsal nerves (B) and misrouted rostral nerves (C) per larva (D-E) Mean swimming covered distance (D) and speed (E). (F) Immunolabelling of sMN axon tracts with Zn-5 antibody in 72-hpf *katna1^+/+^*(n= 57), *katna1^-/-MZ^* (n= 61) and *katna1^-/-MZ^* larvae injected with mouse *TTLL6* (n= 56) or *TTLL11* (n= 54) mRNAs (G-I) Quantifications of dorsal nerve and motility defects of larvae analysed in panel F. (G) Percentage of larvae with dorsal nerve defects. (H) Mean number of split/misguided dorsal nerves per larva. (I) Mean larval swimming speed in the escape-touch response test. Swimming speed values were extracted from tracking analysis of 72-hpf *katna1^+/+^* (n= 65), *katna1^-/-MZ^*(n= 74) and *katna1^-/-MZ^* larvae injected with the mRNAs encoding mouse TTLL6 (n= 78) or TTLL11 (n= 78). (A, F) Full arrowheads and full arrows point at normal rostral and dorsal projections while empty arrowheads and empty arrows indicate misguided rostral and dorsal tracts, respectively. Scale bar: 25 μm. (B-C, G-H) Quantifications were performed on 24 spinal hemisegments located around the yolk tube per larva. (B-E, H-I) Box and Whisker graphs; *p≤0.05; **p≤0.01; ***p≤0.001; ns: non-significant; Kruskal–Wallis ANOVA test with Dunn’s post hoc test. Whiskers indicate the Min/Max values. (G) ***p≤0.001; Chi^2^ test.

### TTLL6 selectively boosts p60-Katanin activity in motor axons to control their targeting

To test whether the rescue of MO^p60Kat/1.3pmol^ morphant phenotypes by TTLL6 overexpression specifically relied on TTLL6-mediated stimulation of p60-Katanin residual activity or whether it involved other MT regulatory proteins (*e.g.*, Spastin), we reproduced the same rescue experiments in the *p60-katanin* null mutant background (*katna1^-/-MZ^*). TTLL6 overexpression (as TTLL11 overexpression) here failed to rescue the axon pathfinding errors and swimming deficit of *katna1^-/-MZ^* mutants (Fig 6F-I). These findings, which strikingly contrasted with the significant impact of TTLL6 overexpression on morphants with residual katanin levels (MO^p60Kat/1.3pmol^), strongly suggest that TTLL6-mediated MT polyglutamylation selectively regulates p60-Katanin activity during motor axon targeting.

### TTLL11 selectively tunes spastin activity in zebrafish navigating motor axons to guide their trajectory

To further characterise the functional selectivity of these long-chain tubulin glutamylases towards MT-severing enzymes *in vivo*, we undertook the same sets of rescue experiments in *spastin* morphant (knockdown) versus mutant (knockout) larvae. In contrast with the results for p60-Katanin, we showed that TTLL11 but not TTLL6 overexpression significantly alleviated the axon pathfinding defects of the rostral motor nerve caused by spastin partial knockdown (injection of 0.2 pmol of MO*sp^ATG^* morpholino; Fig 7A-B). This selective improvement of *spastin* morphant axon phenotypes by TTLL11-mediated polyglutamylation was associated with a slight but significant rescue of larval swimming speed compared to control larvae or to *spastin* morphants overexpressing *TTLL6* (Fig 7A,C-D). Here again, to assess whether the beneficial effect of TTLL11 on *spastin* morphant phenotype relied on TTLL11-driven enhancement of spastin activity or involved other MT-severing proteins, we overexpressed TTLL11 or TTLL6 in the *spastin* CRISPR/Cas9 *null* mutant (*sp^C68X/C68X^*; Jardin et al., 2018). Unsurprisingly, TTLL11 overexpression failed to rescue the rostral motor axon defects of *spastin* null mutants (Fig 7E-G), identifying TTLL11 as a selective regulator of spastin activity in navigating motor axons.

**Figure 7.**
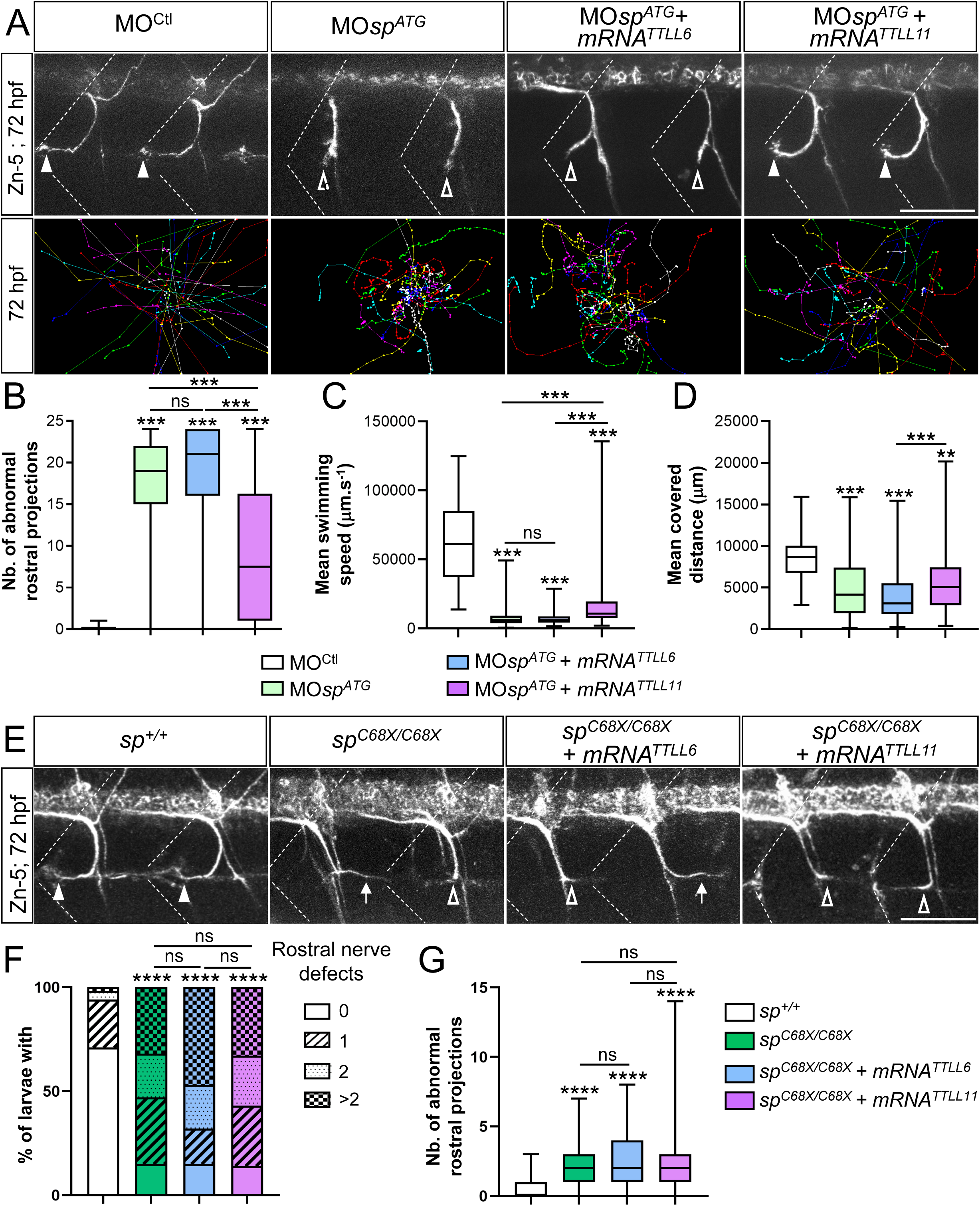
Selective regulation of spastin activity by TTL11 controls zebrafish motor circuit wiring and larval motility. (A) Upper panels: Immunolabelling of sMN axon tracts using Zn-5 antibodies in 72-hpf Tg(*Hb9*:GFP) larvae injected with MO^Ctl^ (n= 69), MO*^spATG^* (n= 79) or co-injected with MO*^spATG^* and mouse *TTLL6* (n= 63) or *TTLL11* (n= 78) mRNAs. Dotted lines delineate lateral myosepta. Lower panels: Tracking analysis of 72-hpf larvae injected with MO^Ctl^ (n= 71), MO*^spATG^* (n= 90) or co-injected with MO*^spATG^* and mouse *TTLL6* (n= 98) or *TTLL11* (n= 124) mRNAs in a touch-evoked escape response test. Each line represents the trajectory of one larva after touch stimulation while the distance between two dots indicates the distance covered by a larva between two consecutive frames. Scale bar: 5 mm. (B-D) Quantifications of the sMN (B) and locomotor defects (C-D) of larvae analysed in panel A. (B) Mean number of abnormal rostral nerves. (C-D) Mean swimming covered speed (C) and covered distance (D). (E) Immunolabelling of sMN axons in 72-hpf *sp^+/+^* (n= 52), *sp^C68X/C68X^* (n= 34) and *sp^C68X/C68X^* larvae injected with mouse *TTLL6* (n= 47) or *TTLL11* (n= 49) transcripts using Zn-5 antibody. (F-G) Quantifications of rostral nerve defects in larvae analysed in panel E. (F) Percentage of larvae with rostral nerve defects. (G) Mean number of abnormal rostral nerves (*i.e.*, defasciculated or missing) per larva. (A, E) Full and empty arrowheads respectively point at normal and defasciculated/missing rostral rostral nerves. Scale bar: 50 μm. (B, F-G) Quantifications were performed on 24 spinal hemisegments located around the yolk tube per larva. (B-D, H) Box and Whisker graphs; **p≤0.01; ***p≤0.001; ns: non-significant; Kruskal–Wallis ANOVA test with Dunn’s post hoc test. Whiskers indicate the Min/Max values. (G) **p≤0.01; ***p≤0.001; Chi^2^ test.

Altogether, our results provide the first *in vivo* proof of concept that selective MT polyglutamylation patterns mediated by specific tubulin glutamylases may rescue the motor neuron and locomotor defects caused by reduced levels of microtubule-severing enzymes *in vivo*.

### TTLL11 selectively rescues the axon degeneration hallmarks caused by spastin haploinsufficiency in a mammalian cellular model of hereditary spastic paraplegia

Importantly, mutations in the *SPG4* gene encoding the MT-severing spastin are responsible for the major form of autosomal dominant hereditary spastic paraplegia (HSP), a degenerative condition of corticospinal tracts (Hazan et al., 1999). While few *SPG4* mutations were shown to have a toxic gain of function in overexpression experiments (Solowska et al., 2014; Qiang et al. 2019), the vast majority of *SPG4* mutations were reported to act through haploinsufficiency (Tarrade et al., 2006; Depienne et al., 2007; Roll-Mecak and Vale, 2008; Kasher et al. 2009; Fassier et al., 2013; Denton et al., 2014; Connell et al., 2016). Thus, increasing spastin activity upon the critical threshold may be a relevant therapeutic strategy in *SPG4*-linked HSP cases.

Our identification of TTLL11 as a selective regulator of spastin activity in zebrafish developing motor neurons prompted us to carry out pilot experiments to assess whether TTLL11 overexpression may alleviate/rescue the axon degeneration hallmarks caused by spastin haploinsufficiency in cultured cortical neurons from spastin heterozygous mutant embryos (Sp+/−; Lopes et al., 2020). We here focused our analysis on axonal swellings, the most significant, easily quantifiable and evolutionarily conserved degenerative hallmark resulting from defective MT dynamics in spastin-deficient neurons (Tarrade et al; 2006; Kasher et al., 2009; Denton et al., 2014; Havlicek et al., 2014).

However, the number of axonal swellings in mouse Sp+/− heterozygous mutant neurons after 6 days in vitro (DIV6) was much lower than that of homozygous mutant neurons, which challenged this rescue analysis (Fig EV4A-B and Tarrade et al., 2006). To overcome this difficulty, we performed a longitudinal analysis of primary cultures of Sp+/+ and Sp+/− cortical neurons and showed that the percentage of axonal swellings significantly increased over time *in vitro* (from DIV6 to DIV9). We showed that DIV9 Sp+/− cortical neurons exhibited a significantly larger number of axonal swellings compared to controls, which was compatible with robust statistical analyses for rescue experiments (Fig EV4A,C). Cultured Sp+/− neurons were thus transduced at DIV2 (before the occurrence of the swellings) with a lentivirus encoding wild-type or catalytically dead (as a negative control) human TTLL11 or mouse TTLL6 glutamylases fused to the GFP, and analysed at DIV9. For each condition, the percentage of swollen axons (Fig 8A-B) and transduced neurons (Fig 8C) were evaluated at DIV9 on fixed neurons immunolabelled with anti-β-III tubulin and anti-GFP antibodies. We showed that wild-type TTLL11 rescued the axonal swelling phenotype caused by spastin haploinsufficiency, unlike TTLL6 or their respective catalytically dead variants (Fig 8A-B), despite similar transduction efficiency (Fig 8C). Notably, TTLL11 overexpression reduced the percentage of axonal swellings in Sp+/− cultures to a level equivalent to that of control neuron cultures (Fig 8B). Importantly, this beneficial effect of TTLL11 on the axonal swelling hallmark of neuronal degeneration was completely lost when the same experiments were conducted in homozygous Sp-/- neurons (Fig 8A-B), confirming that TTLL11 also selectively tunes spastin activity in mammalian cortical neurons. These data provide compelling evidence that enhanced TTLL11-mediated tubulin polyglutamylation prevents (or delays) the axonal phenotypes caused by defective *spastin* gene dosage in animal and cellular models of *SPG4*-linked HSP, revealing an evolutionarily conserved mechanism (Fig 9).

**Figure 8.**
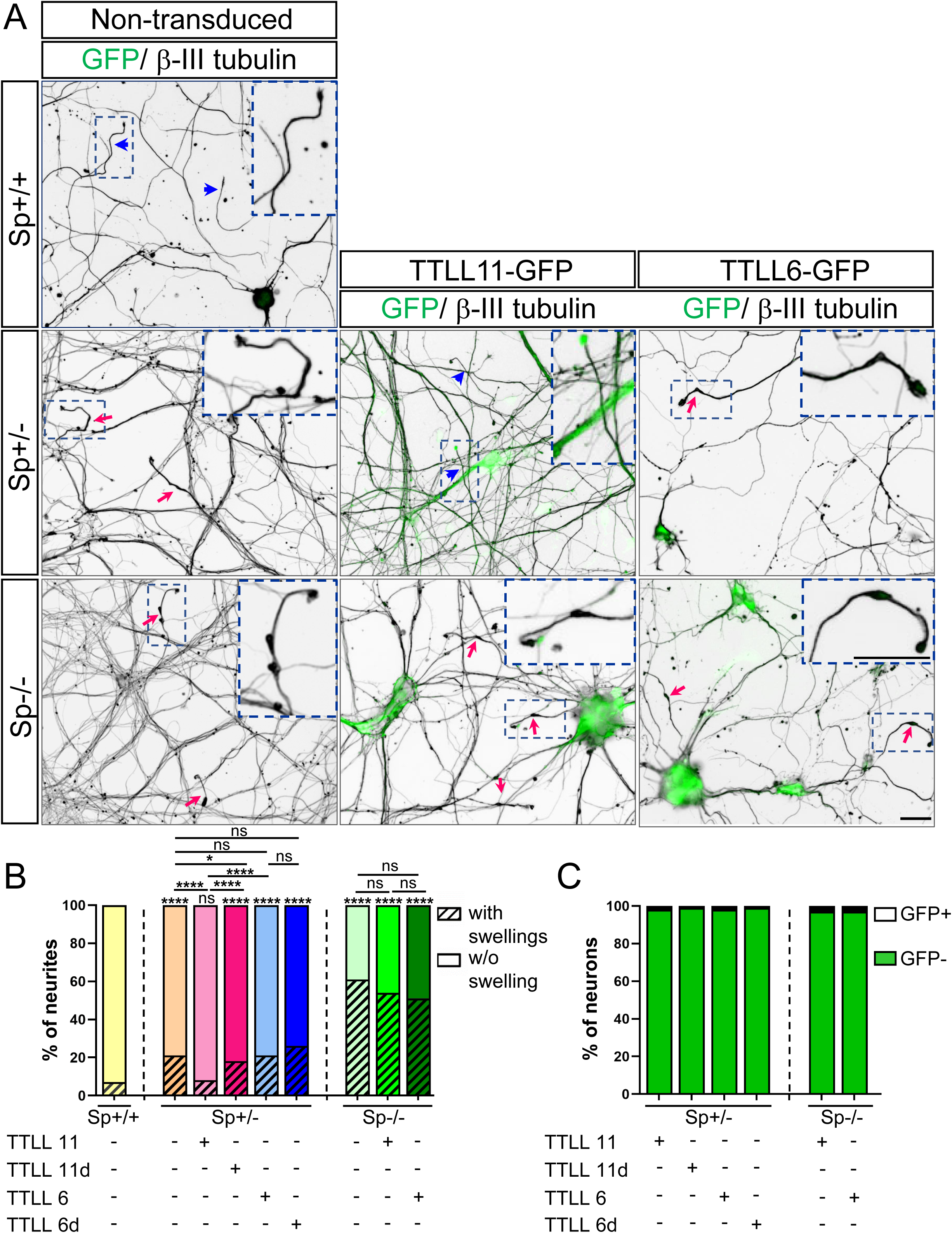
TTLL11, but not TTLL6, rescues the axonal swelling phenotype caused by spastin haploinsufficiency in mammalian cortical neurons. (A) Immunolabelling of β-III tubulin and GFP on DIV-9 Sp+/+, Sp+/− and Sp-/- cortical neurons transduced or not at DIV 2 with lentiviruses encoding wild-type or catalytically dead variants of mouse TTLL11 or TTLL6. Blue arrowheads and pink arrows respectively indicate non-swollen and swollen axons. Insets are higher magnifications of the distal part of the axons framed in the boxed region of each corresponding image. Scale bars: 20μm (B-C) Percentage of swollen axons (B) and transduced neurons (C). Quantifications were performed on at least 400 axons and 500 neuronal somata per condition pooled from two or three independent experiments. *p≤0.05; **p≤0.01; ***p≤ 0.001; Chi2 test.

**Figure 9.**
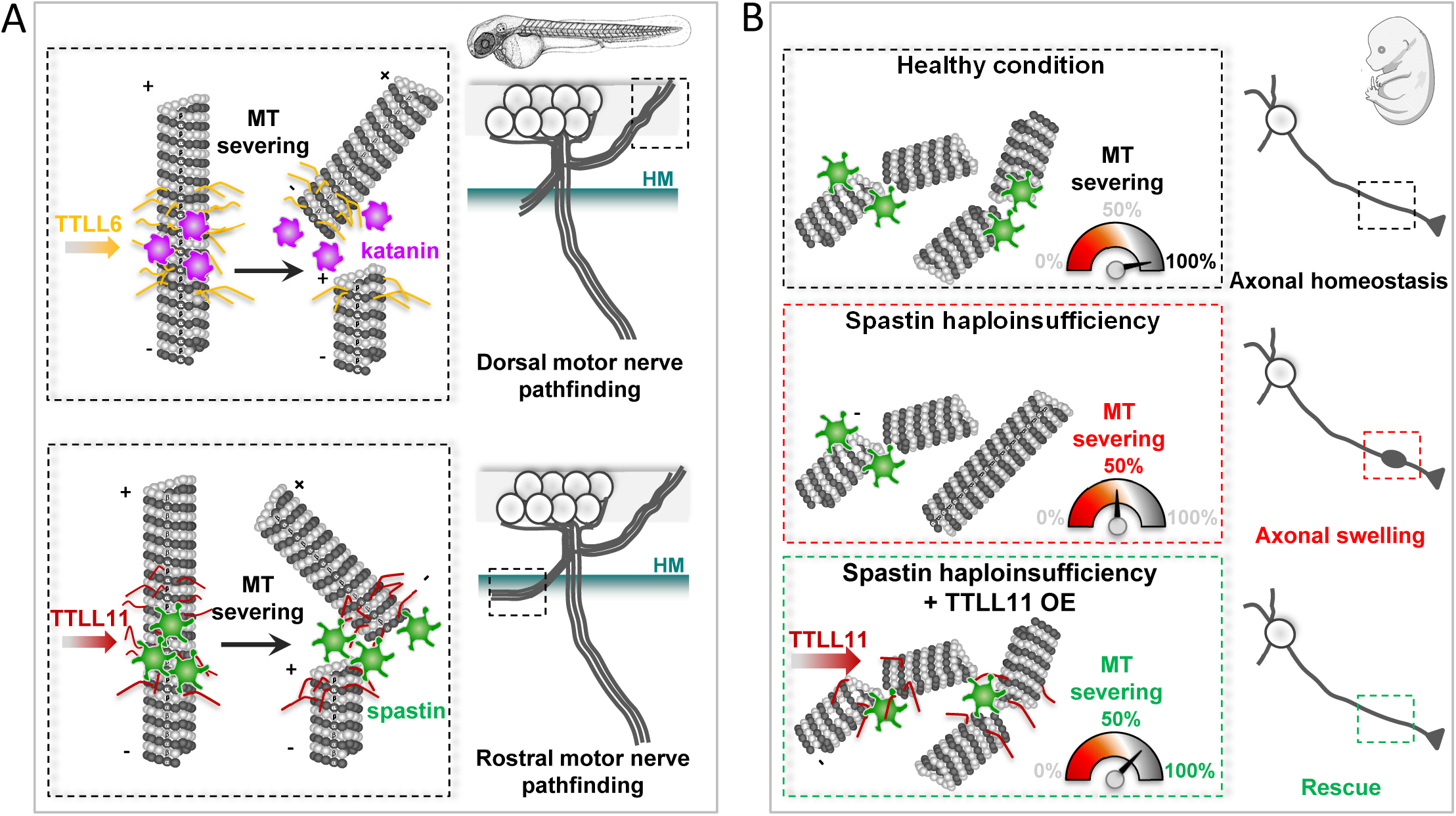
Schematic representation of MT-severing enzyme selective regulation by specific TTLL enzymes in zebrafish and mammalian neurons. (A) Upper panel: TTLL6-mediated tubulin polyglutamylation selectively tunes p60-katanin activity in zebrafish dorsally-projecting secondary motor axons to control their targeting. Lower panel: TTLL11-driven MT polyglutamylation is required for accurate axon pathfinding of rostrally-projecting secondary motor neurons in zebrafish larvae. (B) Spastin haploinsufficiency induces axonal swellings in mammalian cortical neurons (middle panel), which can be selectively rescued by promoting TTLL11-mediated tubulin polyglutamylation, most likely through the boosting of the residual spastin activity upon the critical threshold (i.e., 50%).

## Discussion

Among the myriad of molecules shown to play key roles in axon navigation, special focus has recently been given to MT regulatory proteins due to their prominent roles in integrating and translating extracellular guidance signals into changes in growth-cone mechanical behaviours (Lowery and Van Vactor, 2009; Dent et al., 2011; Kolodkin and Tessier-Lavigne, 2011).

The MT-severing enzyme p60-katanin is known to be a critical player in axon elongation (Ahmad et al., 1999; Karabay et al., 2004; Butler et al., 2010) and branching (Yu et al., 2008). Using loss-of-function and rescue approaches during zebrafish embryogenesis, we here unveil a key role for p60-Katanin in spinal motor axon navigation and targeting. Intriguingly, p60-Katanin regulates this developmental process in a dose-dependent manner, emphasizing the precise fine-tuning of its activity *in vivo*. Interestingly, the orthologue of p60-katanin in *Drosophila* exerts a MT-plus-end depolymerising activity *in vitro* in addition to its known severing activity (Zhang et al., 2011; Diaz-Valencia et al., 2011). The balance between these two functions seems to be influenced by its concentration, with the maximal depolymerisation rate occurring at lower concentrations than the maximal severing rate (Diaz-Valencia et al., 2011). This dual function of p60-katanin was reported to regulate MT dynamics at the actin cortex to fine-tune the migration of *Drosophila* S2 cells by suppressing protrusion. (Zhang et al., 2011). Thus, if applicable *in vivo* in a vertebrate model, this concentration-dependent versatile role of p60-katanin may provide an explanation for the dose-dependent effects of p60-Katanin depletion on axon targeting.

Furthermore, we have here demonstrated that p60-katanin and its related protein spastin have non-overlapping key roles in zebrafish motor axon navigation. This is consistent with their differential regulation of MT remodelling in non-neuronal cells (Lacroix et al., 2010; Shin et al., 2019) and axon branch formation in cultured hippocampal neurons (Yu et al., 2008). The functional specificity of these MT severers in axon branching has so far been linked to their distinct subcellular distribution in neurons and the differential regulation of their activities by the MT-associated protein Tau, which more efficiently protects MTs from being severed by p60-katanin than by spastin (Qiang et al., 2006; Yu et al., 2008). While these mechanisms could also underlie the functional specificity of spastin and p60-katanin in axon navigation, our analysis provides compelling *in vivo* evidence for a central role of MT polyglutamylation in their specificity.

Indeed, we here demonstrate that TTLL6 and TTLL11 selectively tune the respective activities of p60-Katanin and Spastin in developing axons to control their targeting. Notably, the interactions between MTs and some MAPs, including the p60-katanin regulator Tau, are sensitive to tubulin polyglutamylation levels *in vitro* (Boucher et al., 1994; Genova et al., 2023), as shown for spastin and katanin MT-severing activities (Lacroix et al., 2010, Zempel et al., 2013; Valenstein and Roll-Mecak, 2016; Shin et al., 2019, Genova et al., 2023). Overall, the model that emerges from these observations is that MT polyglutamylation patterns generated by distinct polyglutamylases may selectively regulate the activity of MT-severing enzymes by concomitantly (i) promoting their catalytic activity (i.e., increasing their coupling with the tubulin C-terminal tail, hexamerization and ATPase activity; White et al., 2007; Valenstein et al., 2016; Shin et al., 2019; Zehr et al., 2020) and (ii) influencing the MT-binding affinity and/or activity of their regulators, including Tau (Yu et al., 2008) and CRMPs (Ji et al., 2018). Moreover, beyond their molecular specificity, the characteristic axon pathfinding phenotypes caused by the depletion of p60-Katanin or Spastin may also be the readout of their involvement as downstream targets of different axon guidance pathways, as shown for spastin main isoforms (Jardin et al., 2018).

Tubulin PTMs, including glutamylation, have been identified as key mechanisms that generate subpopulations of MTs with distinct properties and potentially specific cellular functionalities (Janke and Magiera, 2020, McKenna et al., 2023). Based on their enzymatic specificity (addition of short or long glutamate side chains) and substrate preference (α-versus β-subunit of tubulin dimers), TTLL enzymes indeed generate different polyglutamylation patterns on MT outer surface (van Dijk et al., 2007). These distinct patterns were shown to fine-tune the activity of MT interactors including MT severers *in vitro* in a selective and gradual manner (Boucher et al., 1994; Lacroix et al., 2010; Valenstein and Roll-Mecak, 2016; Zheng et al., 2022; Genova et al., 2023). However, *in vivo* analyses confirming their crucial role in the selective regulation of MT interactors were still lacking until very recently. Excessive tubulin glutamylation due to carboxypeptidase (CCP) depletion was lately shown to impair axonal transport and cause neurodegeneration in mice and humans (Magiera et al., 2018; Shashi et al., 2018). Interestingly, these axonal transport defects and associated Purkinje cell degeneration caused by CCP1 depletion were shown to be selectively rescued by the knockout of the α-tubulin glutamylase TTLL1 but not by the β-tubulin glutamylase TTLL7 knockout (Bodakuntla et al., 2021). This seminal study thus establishes the relevance of TTLL selectivity towards a specific tubulin isoform in neuronal homeostasis. Adding another level of complexity to the tubulin code, our study shows that two long chain tubulin glutamylases (TTLL6 and TTLL11) with similar substrate specificity (i.e., α-tubulin; van Dijk et al., 2007) are not functionally redundant and selectively boost the activity of key MT-severing enzymes in the wiring of vertebrate neuronal circuits underlying locomotion. Indeed, TTLL6 and TTLL11 may tune a specific MT severer by generating distinct MT glutamylation patterns (i.e., the number of added glutamate residues or the modification site on the tubulin tails). Identifying the glutamylation pattern specific of each TTLL and their impact on prime MT-dependent physiological processes will undoubtedly represent a major technological challenge for the years to come. Furthermore, our data together with previous studies exploring the tubulin tyrosination/detyrosination cycles in the mouse developing brain (Erck et al., 2005; Marcos et al., 2009; Pagnamenta et al., 2019) emphasise the urge to dissect the involvement of tubulin-modifying enzymes in physiological and pathological neuronal circuit wiring.

Finally, the rescue experiments in mammalian cortical neurons from spastin heterozygous mutant mice revealed that the selective capacity of TTLL11 versus TTLL6 to boost residual spastin activity in embryonic neurons is evolutionarily conserved. It further provides the proof of concept that selective modification of tubulin glutamylation pattern, herein through TTLL11 overexpression, rescues the degenerative hallmarks caused by spastin haploinsufficiency in a mouse cellular model of *SPG4*-linked HSP. This finding may pave the way towards the development of innovative MT-based therapeutic strategies in this major form of HSP. Moreover, together with a recent study showing that disrupting tubulin-alpha4a polyglutamylation rescues the key pathological features of a humanised Tauopathy mouse model (Hausrat et al., 2022), our work pinpoints tubulin polyglutamylation as an attractive therapeutic target in MT-related neurodegenerative disorders (Rogowski et al., 2021).

In conclusion, we identify the specific roles of two *in vitro* biochemically undistinguishable tubulin polyglutamylases in axon guidance processes through the selective activation of defined microtubule-severing proteins. This allowed us to show how subtle variations in the tubulin code led to different functional readouts *in vivo*. Our findings also point at enzymes regulating tubulin polyglutamylation, as potential therapeutic targets in pathological conditions associated with reduced levels of MT-severing enzymes.

## Materials and Methods

### Zebrafish maintenance

Zebrafish embryos (*Danio rerio*) were obtained from natural spawning of Tg(*Hb9*:GFP) transgenic fish (Flanagan-Steet et al., 2005), *p60-katanin* mutants *(katna1^sa18250^*; Sanger Centre, purchased from the ZIRC) or spastin CRISPR/Cas9 mutants (Sp*^C68X/C68X^;* Jardin et al., 2018). All embryos were maintained at 28°C in E3 medium (5 mM NaCl/0.17 mM KCl/0.33 mM CaCl_2_/0.33 mM MgSO_4_/0.00001% (w/v) Methylene Blue) and staged by hours post-fertilisation (hpf) and gross morphology according to Kimmel et al. (1995). To prevent pigment formation, 0.2 mM of 1-phenyl-2-thiourea (PTU, Sigma) was added to the E3 media from 24 hpf onwards.

All our experiments were made in agreement with the European Directive 210/63/EU on the protection of animals used for scientific purposes, and the French application ‘Décret 2013-118’. The fish facility has been approved by the French ‘Service for animal protection and health’, with the approval number A-75-05-25.

### Morpholinos and RNA injections

Morpholino oligonucleotides (MO) blocking *p60-katanin/katna1, ttll6, ttll11* or *spastin* translation initiation sites as well as the standard control MO were developed by GeneTools (Philomath, USA) and designed as follows:

MO^ctl^: 5’-CCTCTTACCTCAGTTACAATTTATA-3’,

MO^p60Kat^: 5’-CTCATTGATCTCCCCCAAACTCATC-3’,

MO^katna1aug1^: 5’-CATCCTGTAAGTTAAAGTGGTCAGT-3’ (Butler et al., 2010),

MO^TTLL6^ : 5’-CTGGTGTCCCCATTCTGATCTCTTC-3’ (Pathak et al., 2011) and MO^TTLL11^ : 5’-CGGCTGATTTGTTATCTCATCTAGG-3’,

MO*sp^ATG^*: 5′-GCTGAAACAGCCACCGAAGAAGCC-3′ (Jardin et al., 2018),

COMO*sp^ATG^*: 5′-GCTCAAACACCCAGCGAACAAGGC-3′ (Jardin et al., 2018).

MO^p60Kat^ was injected at 1.3 and 3.4 pmol/embryo, MO^katna1aug1^ at 0.9 pmol/embryo, MO^TTLL6^ at 0.2 pmol/embryo, MO^TTLL11^ at 0.8 pmol/embryo, MO*sp*^ATG^ and COMO*sp^ATG^* at 0.4 or 0.2 pmol/embryo. Universal control MO^ctl^ morpholino was injected at all these doses depending on the controlled knockdown experiment. All morpholinos were injected at the two-cell stage. Human full-coding *KATNA1* cDNA was isolated from a human fetal brain Marathon-ready cDNA collection (Clontech, Invitrogen) with the following forward and reverse primers: 5’-ATATAGGATCCATGTACCCATACGATGTTCCAGATTACGCTAGTCTTCTTATGATTAGTGAG-3’ and 5’-ATATATCTAGATTAGCATGATCCAAACTCAAATATC-3’, and subsequently cloned into pCS2+ BamH1/XbaI restriction sites for rescue experiments. Zebrafish *spastin* full-length cDNA was amplified and HA-tagged by PCR from the IMAGE clone BG728071 with 5’-ATACTCGAGCAAGCTTGATTTAGGTGA-3’ forward and 5’-GGCTCTAGATCAAGCGTAGTCAGGCACGTCGTAAGGGTAACTAGCGCCTACGCCAGTCGTGTCTCCGT-3’ reverse primers, and cloned into pCS2+ using the XhoI/XbaI restriction sites included in the primers. Human *KATNA1*, mouse *TTLL6* and *TTLL11* (cDNAs provided by C Janke), as well as zebrafish *spastin* mRNAs were *in vitro* transcribed from the corresponding linearized pCS2+ constructs using the SP6 mMessage mMachine kit (Ambion) and injected at one-cell stage. For rescue experiments, *KATNA1* mRNA was injected at 120 or 180 pg/embryo, *spastin* transcript at 150 pg/embryo, and both mouse *TTLL6* and *TTLL11* mRNAs at 120 pg/embryo.

### Genomic DNA isolation and genotyping

Genomic DNA was isolated by incubating larval heads 2 hours at 55°C in lysis buffer (100mM Tris Hcl/2mM EDTA pH8/0.2% Triton X-100/250 μg/mL proteinase K). Homozygous and heterozygous *katna1*mutants were identified by PCR amplification followed by DNA sequencing (GENEWIZ). The primers used for genotyping were the following:

Katna1sa18250_FOR: 5’-GTAGTACGGAAATCCTCTGTCC-3’

Katna1sa18250_REV: 5’-TTGCTTTGATCTAAGAAACCGG-3’.

*Spastin* (*sp^C68X/C68X^*) mutants were genotyped as previously described in Jardin et al. (2018).

### RNA extraction and RT-qPCR analysis

For sequence analysis of *katna1* mRNA, total mRNA of 24-hpf *katna1^+/+^* and maternal zygotic *katna1^-/-MZ^* zebrafish embryos were extracted with Trizol according to the manufacturer’s instructions. *Katna1* transcripts were reverse-transcribed and PCR-amplified using the SuperScript™ III One-Step RT-PCR System with Platinum® Taq (ThermoFisher Scientific, France) from 200 ng of *katna1^+/+^* and *katna1^-/-MZ^* RNA extracts using the following primers:

Katna1RTPCRex3-4_FOR: 5’-ATGTGGAGCACAGATCGTCTCCATGTG-3’

Katna1RTPCRex5-6_REV: 5’-CAGCAATGTCATCCCATGTGACATTGGG-3’.

For RT-qPCR analysis of *katna1* homologous gene expression levels, RNAs were extracted from 5 independent pools of 10 *katna1^+/+^*, 10 *katna1^-/-MZ^* and one pool of 10 wild-type embryos (at 24 hpf) using the RNeasy Mini-kit (Qiagen, France) and reverse-transcribed using the Superscript RT II Kit with random hexamers (Invitrogen, ThermoFisher Scientific, France). qPCR was performed using a SYBR Green master mix (EurobioGreen qPCR Mix, Hi-ROX; Eurobio Scientific, France) using the primers listed in Table 1. The relative quantification method was used to calculate the expression levels of the genes of interest normalized to *lsmb12* and relative to the cDNAs from 24-hpf wild-type embryos. Values are shown as means ± SEM.

**Table 1:**
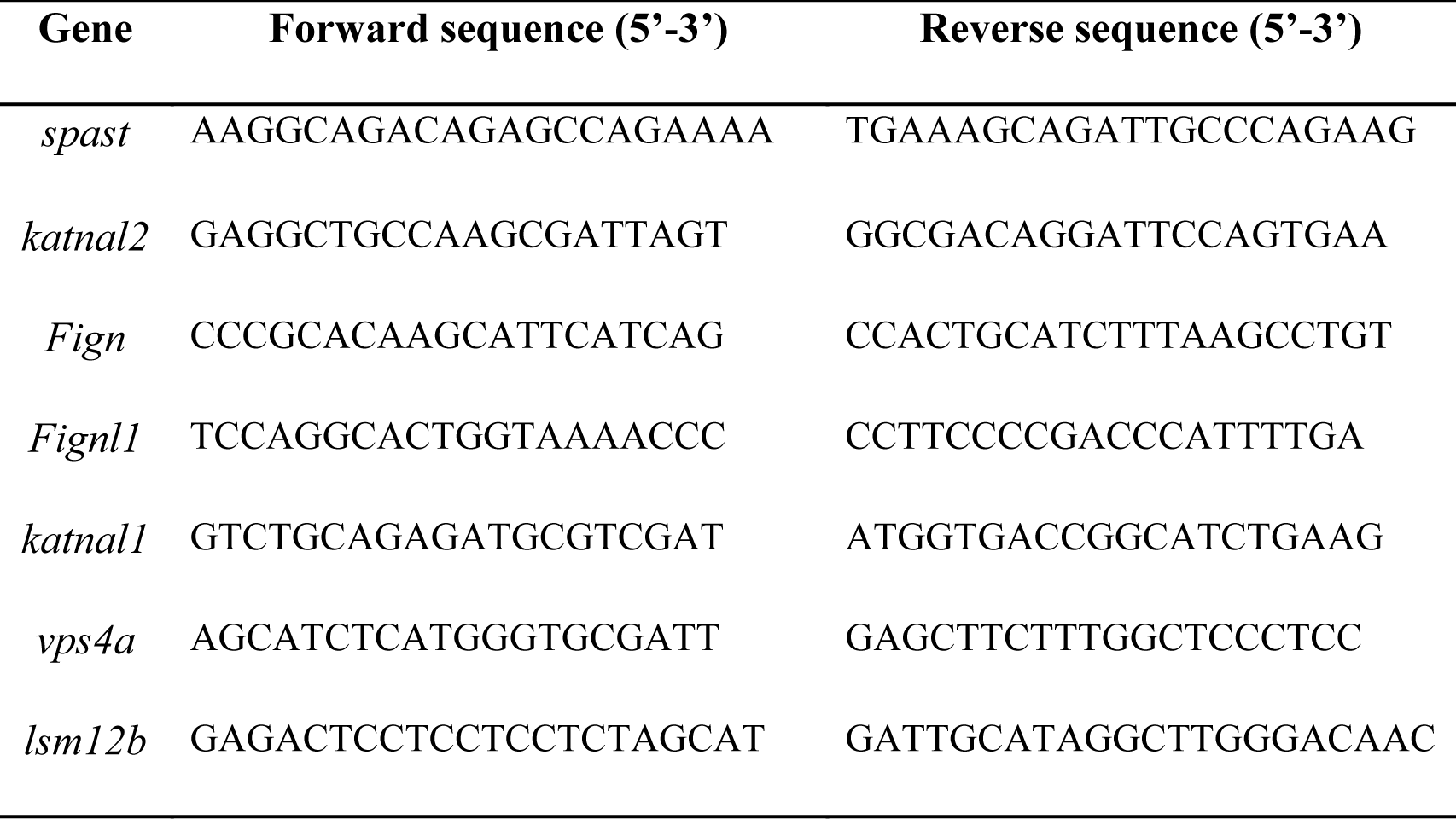
Primers used for RT-qPCR analysis.

### Whole-mount immunohistochemistry

Zebrafish embryos were fixed in PBS/4% paraformaldehyde during 2 hours at room temperature, washed 3 times with PBT1% (1% Triton X-100 in PBS) and permeabilised using a 0.25% trypsin solution (Gibco) at 25°C when embryos were older than 24 hpf. Embryos were then blocked for two hours in PBT1% supplemented with 10% normal goat serum and incubated with primary antibodies overnight at 4°C in PBT1%/1% normal goat serum. The following primary antibodies were used: Zn-5 (1/200; ZIRC, University of Oregon) and GFP (1/1000; Molecular Probes). After several washes in PBT1%, embryos were incubated overnight at 4°C with the appropriate secondary antibody (Alexa Fluor 488 or 555 at 1/1000, Molecular Probes).

For glutamylated Tubulin staining, embryos were fixed in Dent’s fixative (80% Methanol/20% DMSO) overnight at 4°C. After rehydration in regressive methanol/PBS-0.5% Tween 20®, embryos were blocked in PBT1% supplemented with 1% DMSO, 1% bovine serum albumin and 5% normal goat serum, and sequentially incubated with primary antibodies, GT335 (1:1000, Adipogen) and PolyE (1:1200; provided by C. Janke), and adequate secondary antibodies. Images were acquired using a fluorescence microscope equipped with an Apotome module (Zeiss, Axiovert 200M), a 20x objective (NA 0.5), the AxioCam MRm camera (Zeiss) and the Axiovision software (Zeiss). Images were processed with the NIH Image J software. Each figure panel corresponds to a projection image from a z-stack of 2-μm sections.

### Time-lapse videomicroscopy assay

Control and MO^p60Kat^-injected Tg(*Hb9*:GFP) embryos were anesthetised with E3 medium containing tricaine and embedded in 0.8% low-melting agarose in a 35-mm glass dish (Iwaki). Time-lapse videomicroscopy recording of spinal motor axon outgrowth was carried out at 28°C in E3 medium (supplemented with tricaine) using a Leica DMI 6000B inverted spinning-disk microscope with a 40x immersion objective (NA 1.4) and a 491-nm 100-mW Cobolt calypso laser over 30 hours. Embryos immobilised at 22 hpf were filmed with a 20x immersion objective over 48 hours. In both cases, z-stacks of 80-μm-thick sections were acquired every 8 minutes with a step size of 1 μm using an EMCCD camera (Photometrics Quantem 512 SC) and the Metamorph software (Molecular Devices) and compiled into time-lapse movies.

### Touch-evoked escape response test and manual tracking

To assess the motor behaviour of control, mutant, morphant and rescued 72-hpf larvae, we performed a touch response test by applying a tactile stimulus with a pair of forceps and analysing the larval escape behaviour under a Leica M165 C binocular stereomicroscope equipped with a Leica IC80 HD camera. Swimming speed and covered distance of each larva were quantified using the Manual Tracking plugin included in the Fiji software as reported in Fassier et al. (2010).

### Whole-mount *in situ* hybridization

A 3’-fragment of *katna1* cDNA was isolated from a collection of 24-hpf zebrafish embryo mRNAs using the SuperScript® III One-Step RT-PCR system (Invitrogen) with the following forward and reverse primers: 5’-AGAGTGGATTTACTCAAGATCAACC-3’ and 5’-AAGCTTGACTTGTACGCAGTGAACC-3’. The RT-PCR product was cloned into the TOPO® TA cloning pcr4 vector (Invitrogen) and sequenced. The *katna1* digoxigenin-labelled sense and antisense riboprobes were synthetized from the linearised recombinant TOPO® TA cloning vector using T7 and T3 RNA Polymerase (Promega) according to the supplier’s instructions. Whole-mount *in situ* hybridisation experiments were performed at 18 and 24 hpf using standard procedures (Macdonald et al., 1994). Pictures were acquired with a binocular stereomicroscope (Leica M165C) combined with an HD camera (Leica IC80 HD), then adjusted for brightness and contrast with the NIH Image J software.

### Primary cultures of cortical neurons, lentiviral infection and immunolabelling

Primary cultures of cortical neurons were prepared as described in Tarrade et al. (2006) from Sp+/+, Sp+/− and Sp-/- mice (provided by the Kneussel lab; Lopes et al., 2020) at E14.5 day post coïtum. Cortical neurons were plated at 600,000 neurons per 35mm petri dishes and maintained for at least 9 days at 37°C in 5% CO_2_. One third of the medium was changed every three days. Cortical neurons were fixed at 6, 7 and 9 days with 4%PFA in 4% sucrose/PBS for 15 minutes at 37°, permeabilized with 0.1% Triton-100 in PBS for 5 min, blocked in 3% BSA/ 5% normal goat serum in PBS for 1h and incubated overnight at 4°C with primary antibodies (β-III tubulin/anti-tuj-1, 1/1000, Biolegend n°801202; anti-GFP, 1/1000, Torrey Pines Biolabs n°TP-401) diluted in the blocking solution. After several washes in PBS, the primary neurons were incubated with secondary antibodies (anti-mouse-568, 1/1000, Molecular probes A11019; anti-rabbit-488, 1/1000, Molecular probes A11008), sir-actin (1/2000, Cytoskeleton, n°CY-SC001) and DAPI for 1h at RT. Neurons were then rinsed in PBS, mounted using the ProLong Gold mounting medium (Invitrogen n° P36934) and imaged using an epifluorescent microscope (DM6000, Leica) with the 40x objective (N.A: 0.5). The number of neurite swellings per 100 nuclei was determined for each condition. For rescue experiments, Sp+/− and Sp-/- cortical neurons were treated at DIV2 (Day In Vitro) with lentivirus suspensions encoding WT or catalytically dead mouse TTLL6 or human TTLL11. Viruses were produced as described previously (Bodakuntla et al., 2020). Primary cortical neurons were transduced with a volume of viral suspension able to transduce more than 97% of neurons. This volume was determined for each lentiviral construct by infecting the primary neurons with serial dilutions of the supernatant containing the viral particles. Cells were fixed at DIV9 and immunolabelled as described above for the non-transduced cells. To ease the identification of axonal swellings in the dense neural network at DIV 9, the number of swollen axons was estimated in at least 100 field of view acquired at the 40x objective (N.A: 0.75) and related to the total number of analysed axons. Each set of experiments was reproduced three times independently. Animal care and use were performed in accordance with the recommendations of the European Community (2010/63/UE) for the care and use of laboratory animals. Experimental procedures were specifically approved by the Ethics committee of the Institut Curie CEEA-IC #118 (authorization #37315-2022051117455434 v2 given by National Authority) in compliance with the international guidelines.

### Statistical Analysis

All data were obtained from at least two to three independent experiments. Statistical significance of the data was evaluated using the non-parametric Mann-Whitney test when comparing two groups assuming non-Gaussian distributions. The Kruskal–Wallis ANOVA test with Dunn’s post-test or the One-Way ANOVA test with Bonferroni’s post-test were used when comparing more than two groups assuming non-Gaussian or Gaussian distribution, respectively. The *Chi^2^*test was used to compare the distribution of the zebrafish motor neuron defects in different experimental conditions. Data distribution was tested for normality using the D’Agostino and Pearson normality test. All Statistical analyses were performed using GraphPad Prism 9.00 (GraphPad Software, San Diego, CA).

## Author Contribution

JH and CF designed the zebrafish part of the project and interpreted the data with DTM and NJ. DTM performed the functional analysis of p60-Katanin, TTLL6 and TTLL11 during zebrafish motor circuit development with the exception of live video-microscopy experiments, which were carried out by CF. CF and JV performed all the zebrafish experiments using the *katna1* mutants as well as *spastin* mutant and morphants with the assistance of CH for RT-qPCR. NJ investigated the differential role of Spastin and p60-Katanin in spinal motor axon targeting with FG. LG provided technical assistance and developed various constructs. CF and JV are members of XN/CF lab and benefited from XN experience in axon guidance processes. CJ and VH performed TTLL cloning. The mouse experiments in spastin knockout mice, generated by MK, were designed and interpreted by CF, MM and CJ. CF and MM performed cortical neuron experiments with the assistance of VH and LL for mice genotyping and lentivirus production. FG, CJ and MM provided insightful comments throughout the study. JH and CF wrote the manuscript, with the editing of MM and CJ.

## Expanded View figure Legends

**Figure EV1.**
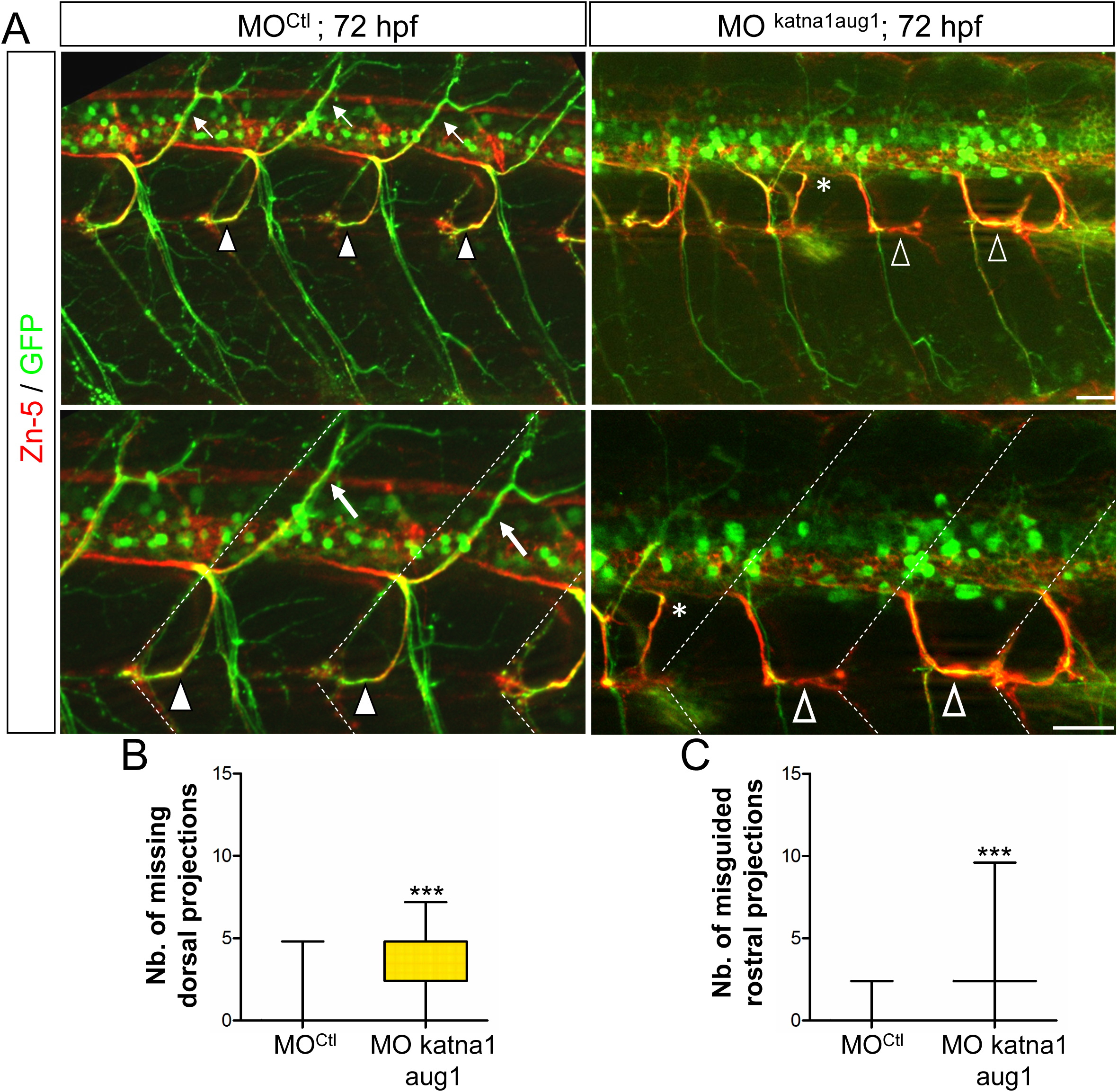
*p60-katanin* knockdown with MO^katna1aug1^ morpholino induces similar spinal motor axon defects to MO*^p60Kat^*morpholino. (A) Immunolabelling of secondary motoneuron (sMN) axon tracts in 72 hours-post-fertilisation (hpf) transgenic Tg(*Hb9*:GFP) larvae injected with control (n=40) or MO^katna1aug1^ (n=40) morpholinos, using Zn-5 and GFP antibodies. Lateral views of the trunk, anterior to the left. Bottom panels represent higher magnifications of top panels. Dotted lines delineate lateral myosepta. Full arrowheads and full arrows point at normal rostral and dorsal nerves, respectively. Empty arrowheads show misguided rostral nerves. Asterisks indicate ectopic sorting points of sMN axons from the spinal cord. (B-C) Quantifications of sMN defects in larvae analysed in panel A. Mean number of missing dorsal nerves (B) and misguided rostral nerves (C) per larva. Quantifications were performed on 24 spinal hemisegments located around the yolk tube per larva. Box and Whisker graphs. ***p≤0.001; Mann Whitney test. Whiskers indicate the Min/Max values.

**Figure EV2.**
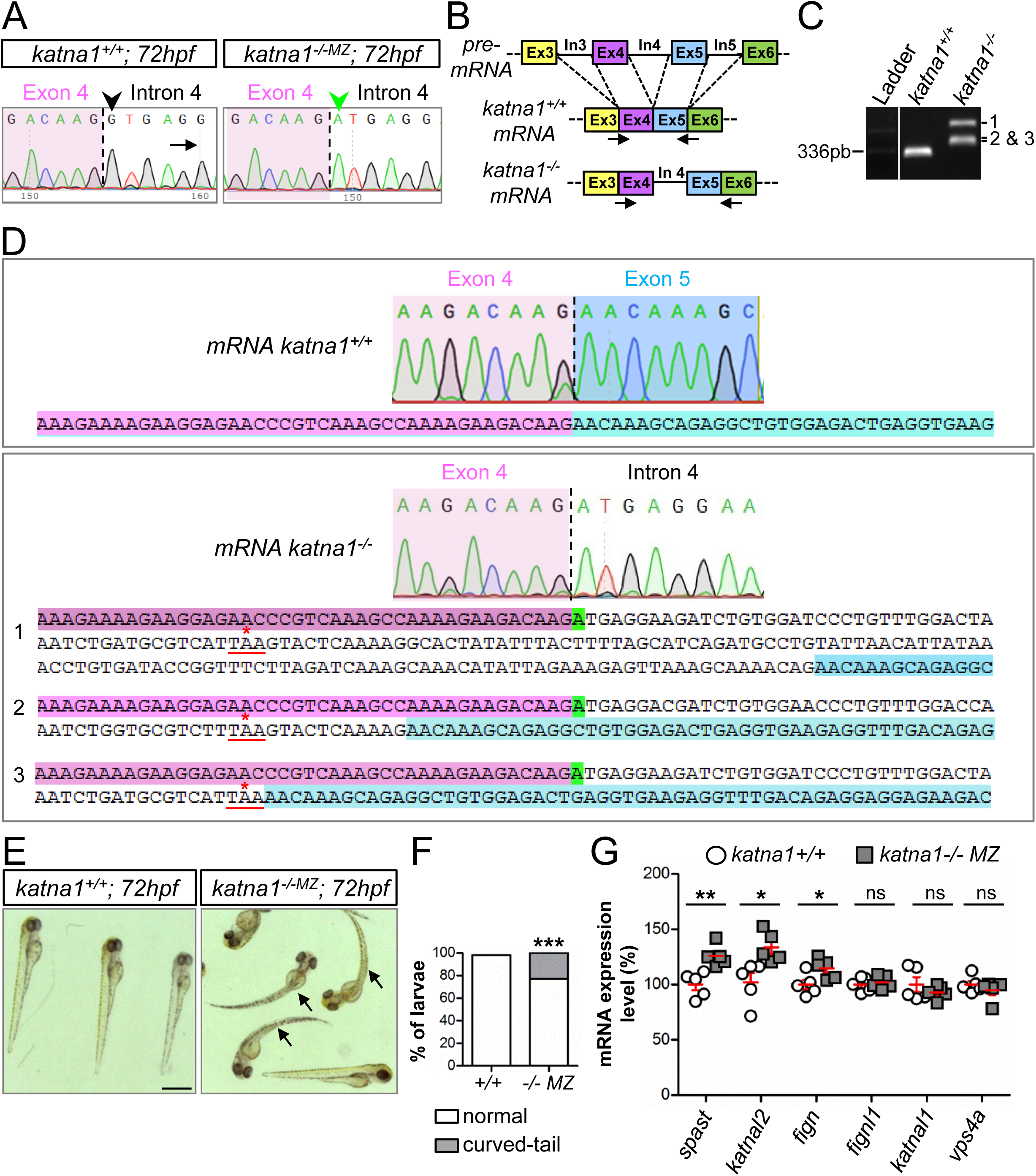
Molecular and morphological characterisation of zebrafish *katna1* mutants. (A) Sequence analysis of control and *katna1* mutant genomic DNA. Dotted line indicates the junction between exon 4 (pink background) and intron 4 (white background). The green arrowhead points at the nucleotide substitution (G>A) affecting the donor splice site of intron 4 (G in control, black arrowhead) in *katna1* mutants. (B) Schematic representation of the RT-PCR strategy used to test the impact of the *katna1* splice-site mutation on *katna1* mRNA splicing. Dotted lines indicate intron splicing. Arrows represent the primers used for RT-PCR analysis. Primers were designed on exon/intron junctions to avoid contamination by genomic DNA amplification. In: intron; Ex: exon. (C) RT-PCR analysis of *katna1* intron-4 splicing on total RNA extracts from *katna1^+/+^* and *katna1^-/-MZ^* maternal zygotic mutant embryos. Homozygous *katna1^-/-MZ^* embryos lack wild-type transcript and show different populations of misspliced transcripts (1, 2 and 3). (D) Sequence analysis of *katna1* misspliced transcripts. Sequences corresponding to exon 4, intron 4 and exon 5 are respectively indicated in pink, white and blue. The splice-site mutation is highlighted in green. Misspliced transcripts include various sized insertions of intron 4, which all lead to a frameshift and the occurrence of a premature stop codon at the same amino-acid position (red asterisk). (E) Gross morphology of 72-hpf control (*katna1^+/+^*) and maternal zygotic *katna1* mutant (*katna1^-/-MZ^)* larvae. Arrows point at the curved-tail phenotype of some mutant larvae. Scale bar: 0.5 mm. (F) Percentage of larvae exhibiting a curved-tail phenotype. ***p≤0.001; Chi^2^ test. (G) RT-qPCR analysis showing the differential expression levels of *p60-katanin*-related genes from the ATPase Meiotic Clade in control and *katna1^-/-MZ^*larvae. * p≤0.05; **p≤0.01; ns: non-significant; Unpaired t-test.

**Figure EV3.**
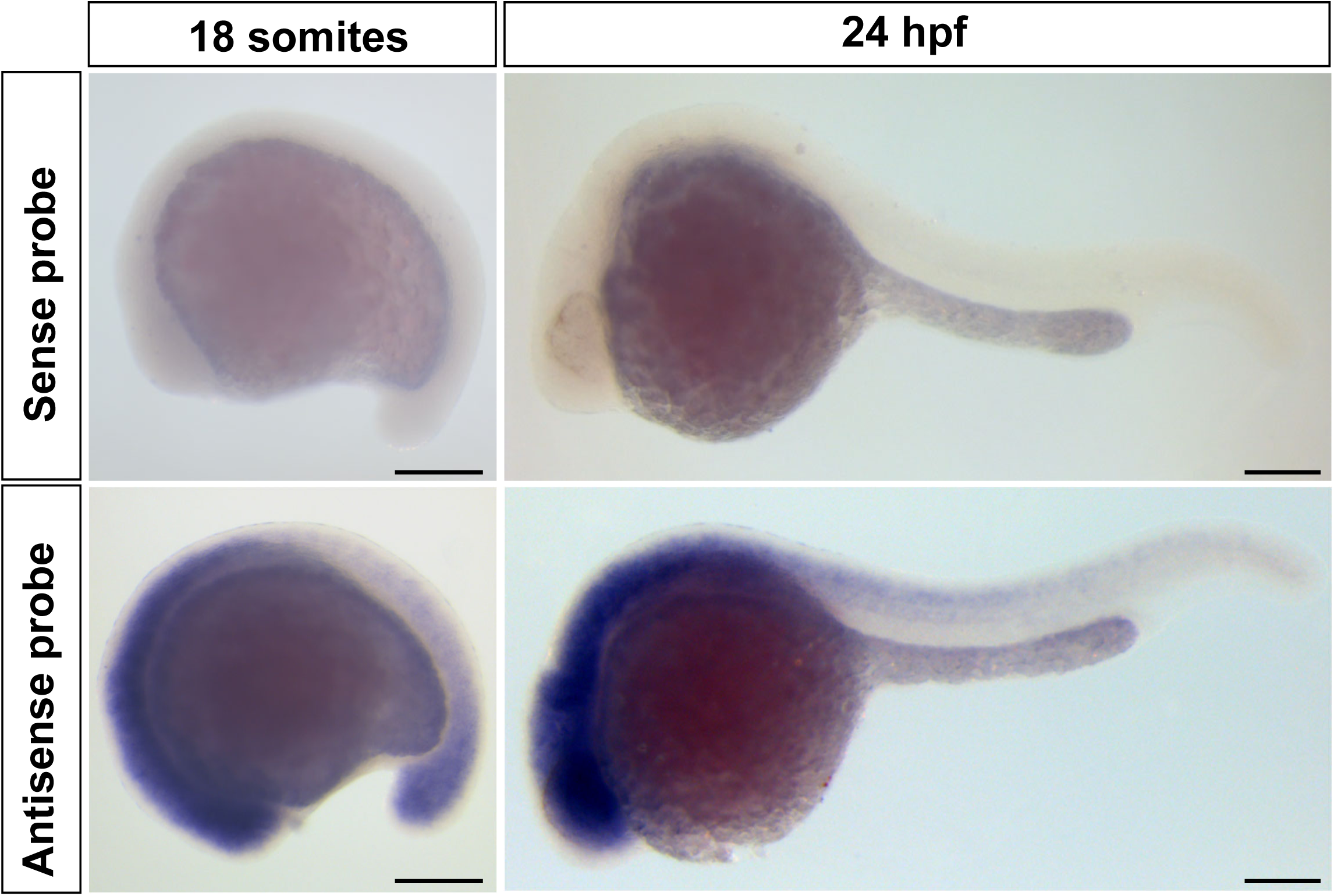
*p60-katanin* transcript is enriched in the zebrafish developing nervous system. Whole-mount *in situ* hybridisation with *p60-katanin* sense (upper panel) and antisense (bottom panel) riboprobes at 18 somites and 24 hours post-fertilisation (hpf). Lateral views of the embryos, anterior to the left. *P60-katanin* is highly enriched in the developing nervous system at both 18 somites and 24 hpf, two stages at which the axons of primary (pMN) and secondary (sMN) motor neurons exit the spinal cord to navigate towards their muscle targets. Scale bars: 200 µm.

**Figure EV4:**
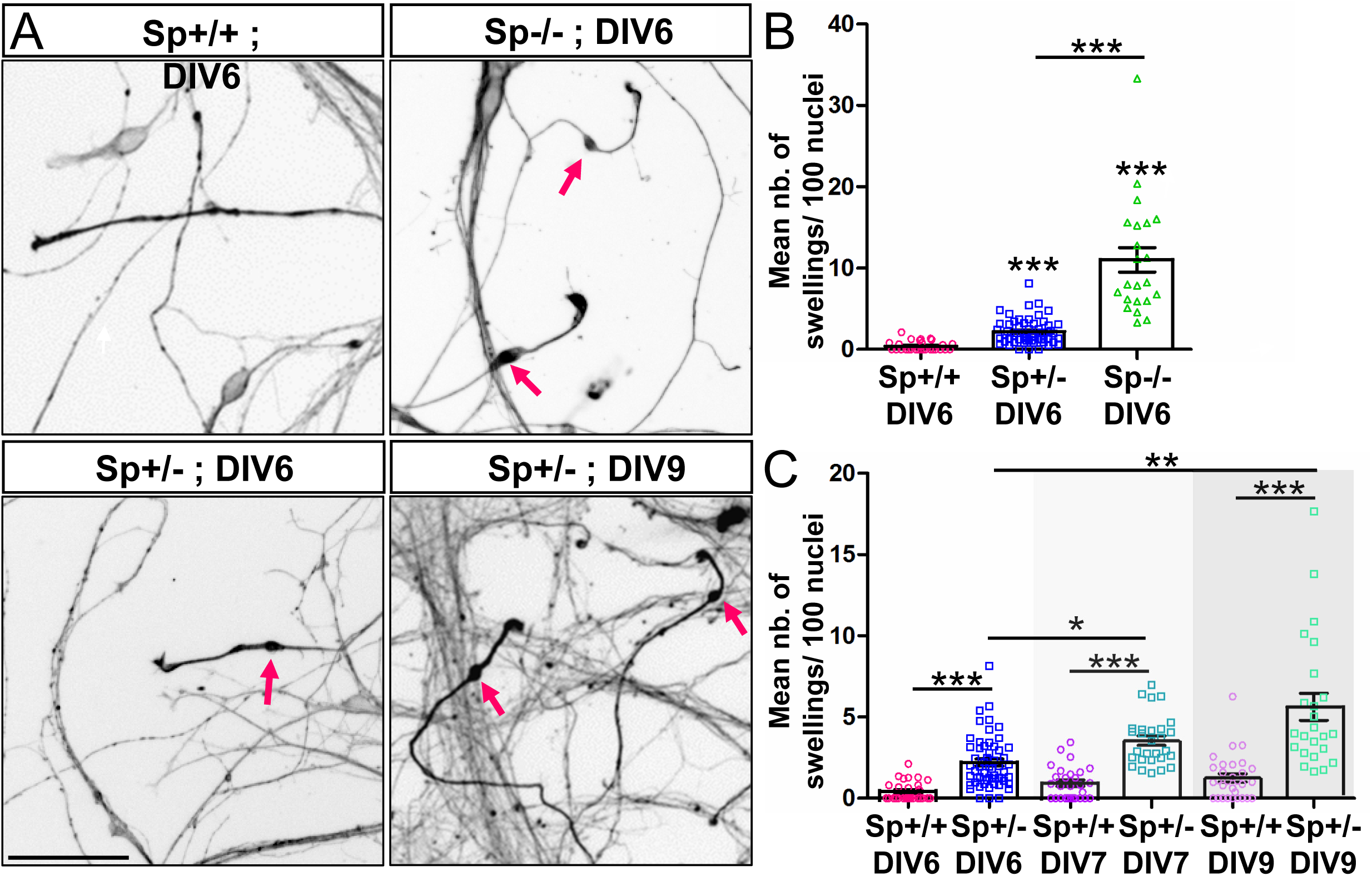
Mouse Sp+/− cortical neurons exhibit a significant number of axonal swellings. (A) Mouse Sp+/+, Sp+/− and Sp-/- cultured cortical neurons immunolabelled with a βIII- tubulin antibody at different days in vitro (DIV). Arrows point at axonal swellings. Scale bars: 50µm. (**B-C**) Mean number of axonal swellings per 100 nuclei. At least 2500 neurons were analysed per condition. *p≤0.05; **p≤0.01; ***p≤ 0.001; Kruskal–Wallis ANOVA test with Dunn’s post hoc test. Error bars are SEM.

## Expanded view movies

**Movies EV1-2: *p60-katanin* knockdown leads to the abnormal splitting of the dorsal nerve.**

**Movie EV1: Time-lapse videomicroscopy of dorsal motor nerve outgrowth in control Tg(*Hb9*:GFP) larvae.** Frames were acquired with a spinning disk microscope and a 40x objective every 8 minutes from 40 to 72 hpf.

**Movie EV2: Time-lapse videomicroscopy of dorsal motor nerve outgrowth in MO^p60Kat^- injected Tg(*Hb9*:GFP) larvae.** Frames were acquired with a spinning disk microscope and a 40x objective every 8 minutes from 40 to 72 hpf.

**Movies EV3-4: *p60-katanin* knockdown strikingly impairs spinal motor axon navigation**

**Movie EV3: Time-lapse videomicroscopy of motor axon outgrowth in control Tg(*Hb9*:GFP) larvae.** Frames were acquired with a spinning disk microscope and a 20x objective every 8 minutes from 20 to 72 hpf.

**Movie EV4: Time-lapse videomicroscopy of motor axon outgrowth in MO^p60Kat^-injected Tg(*Hb9*:GFP) larvae.** Frames were acquired with a spinning disk microscope and a 20x objective every 8 minutes from 20 to 72 hpf.

**Movies EV5-10: The swimming deficit of *p60-katanin* morphants is rescued by overexpression of human *KATNA1* transcript but not by zebrafish *spastin* mRNA.**

**Movie 5: Touch-evoked mobility of 72-hpf control larvae.**

**Movie 6: Touch-evoked mobility of 72-hpf MO^p60Kat*/*1.3pmol^-injected larvae.**

**Movie 7: Touch-evoked mobility of 72-hpf larvae injected with both MO^p60Kat*/*1.3pmol^ and 120 pg of human *KATNA1*mRNA.**

**Movie 8: Touch-evoked mobility of 72-hpf MO^p60Kat*/*3.4pmol^-injected larvae.**

**Movie 9: Touch-evoked mobility of 72-hpf larvae injected with both MO^p60Kat*/*3.4pmol^ and 180 pg of human *KATNA1* mRNA.**

